# Metagenome-assembled genomes reveal water-mass-associated distribution ecotypes of marine Archaea in the Western Arctic Ocean

**DOI:** 10.64898/2026.11.02.763590

**Authors:** Kari Green, R. Eric Collins

## Abstract

Archaea are important contributors to marine biogeochemical cycling, yet their ecology remains poorly characterized in the Arctic Ocean. The three dominant marine archaeal lineages, Marine Groups I, II, and III (MG-I, MG-II, and MG-III), are generally reported to have distinct, clade-consistent vertical distributions that are attributed to light penetration and temperature. Here, we used shotgun metagenomic sequencing of seawater collected during the 2015 U.S. Arctic GEOTRACES expedition (GN01) to recover archaeal metagenome-assembled genomes (MAGs) from the Western Arctic Ocean and to relate their relative abundances to water mass structure. Temperature, salinity, silicate, and N*^*^* distinguished four water mass layers: the summer surface layer, the Polar Mixed Layer, Atlantic Water, and Arctic Ocean basin water. Of 155 MAGs recovered from a metagenomic co-assembly, 17 were archaeal and represented all three marine groups (four MG-I, eleven MG-II, and two MG-III). Abundance-based clustering of samples from two deep central Arctic stations separated microbial communities according to water mass. Mapping MAG relative abundance across several hundred samples revealed surface, mid-depth, and deep distribution ecotypes; members of every marine group displayed more than one ecotype, and ecotype was not consistent with MAG phylogeny. These results indicate that archaeal distributions in the Western Arctic Ocean are more closely tied to water mass stratification than to clade identity, and that previously reported clade-specific depth distributions of marine Archaea may partly reflect the limited resolution of amplicon-based surveys and shallow sampling.

## 1 Introduction

The Arctic Ocean is an environment characterized by habitats with extremes in light, temperature, and salinity (Collins et al., 2010; Junge et al., 2001). Globally, the Arctic Ocean is small, accounting for *∼*3% of the world’s oceans by area (Chang and Devol, 2009; Jensen et al., 2019); however, it contains *∼*20% of the world’s continental shelf (5.1 *×* 10^6^ km^2^) (Chang and Devol, 2009). This ocean is semi-enclosed by land with approximately 50% of its surface area (9.5 *×* 10^12^ m^2^) overlying continental shelves (Jensen et al., 2019). The Arctic Ocean consists of two major basins, the Eurasian and the Amerasian Basins, separated by the Lomonosov Ridge (Rudels, 2015). These larger basins can be subdivided by bathymetric regions called subbasins. The Eurasian Basin is further branched into the Nansen Basin and Amundsen Basin (the deepest, *∼*4500 m), by the Gakkel Ridge, while the Western Amerasian Basin is comprised of the Canada and Makarov subbasins separated by the Alpha and Mendeleev Ridges (Jensen et al., 2019; Rudels, 2015). Water circulation into the Arctic Ocean occurs in both the east, where Atlantic water enters through the deep passage (2600 m) of the Fram Strait (Rudels, 2015), and the west, where northern Pacific water enters through the narrow and shallow (50 m) Bering Strait (Jensen et al., 2019; Rudels, 2015). Atlantic water moves through the Fram Strait and Barents Sea via the Nordic Sea and through the Canadian Archipelago by Baffin Bay and Davis Strait (Rudels, 2015). Fram Strait is the only passage that allows two-way movement of water into and out of the Arctic Ocean. Upon entering the Arctic Ocean, the warm Atlantic water encounters and melts sea ice. In the uppermost layer, the water is cooled and homogenized as ice forms through brine rejection and thermohaline convection (Rudels, 2015). In the lower layers, the Atlantic water cools and sinks (Whitmore et al., 2019). Pacific water entering the Chukchi Sea through Bering Strait undergoes biochemical and physical changes, such as surface heating and cooling, and internal mixing, during its transit over the shelves and into the Arctic Ocean Basins (Gong and Pickart, 2016). The western region is strongly defined by its thermocline and macronutrient characteristics. Surface water circulation in the Amerasian Basin is strongly influenced by the anticyclonic Beaufort Gyre and the Transpolar Drift, a strong current that advects water from the Chukchi, Eastern Siberian, and Laptev Seas across the central Arctic Ocean (Chang and Devol, 2009; Gong and Pickart, 2016; Whitmore et al., 2019, 2022). Generally, the Arctic Ocean can be separated vertically into three major layers: the polar surface water (PSW), Atlantic water, and deep bottom waters, although variations may be observed within the different basins. In the Amerasian Basin, surface waters, commonly referred to as the Polar Mixed Layer (PML) of the PSW (Rudels, 2015), originate primarily from Pacific Ocean water that is advected through the Bering Strait (Jensen et al., 2019). The PML is fresher and colder and can extend variably 0-50 m (Farmer et al., 2021; Jensen et al., 2019). The stratification observed here is strong enough to reduce the amount of mixing with lower layers which allows the low salinity water in the PML to cool enough for ice to form in the winter. This ice layer persists throughout the year in the central Arctic Ocean (Rudels, 2015). Ice formation allows the PSW to become homogenized in winter through freezing, brine release, and thermohaline convection processes. In summer, this ice will melt resulting in re-stratification, which again is removed during freezing in the fall and winter.

Below the PML, salinity sharply increases through the halocline with temperatures remaining close to freezing. Pacific water entering the Bering Strait, less saline than the Atlantic water, but more saline than the PML, forms an intermediate layer. Summer Bering Sea water sits above, and winter Bering Sea water sits below (Rudels, 2015) and mainly remains within the Amerasian Basin. The water below the halocline and intermediate layer is known as the Atlantic layer coming in from Fram Strait. This water enters the Arctic Ocean and travels in a counterclockwise motion towards Svalbard (Fram Strait Branch; FSB) and further eastward toward the Barents Sea (Barents Sea Branch; BSB) becoming colder and more saline (Jensen et al., 2019). As the FSB travels, it mixes with the colder, less saline water above the halocline and continues to cool as it moves into the Makarov and Canada Basins (Jensen et al., 2019; Woodgate et al., 2001). Within these Basins, the FSB can typically be found between 350 and 800 m. The BSB lies below the FSB and extends to 1150 m. Water below 1200 m originates from the far North Atlantic (Greenland and Norwegian Seas) both entering and exiting through the Fram Strait (Jensen et al., 2019). The residence times in the Makarov and Canada basins differ (300 and 400 years, respectively) due to the separating Alpha-Mendeleev Ridge (Jensen et al., 2019). Although distinct, these basins are still subject to mixing. The Arctic Ocean is highly climate relevant due to its impact on the formation of North Atlantic Deep Water, a driver of the Atlantic Meridional Overturning Circulation, as well as the albedo effect it provides due to high levels of ice coverage (Whitmore et al., 2022). In recent years we have witnessed the loss of sea ice, particularly multi-year sea ice due to increased Arctic warming (Box et al., 2019; Chang and Devol, 2009; Edwards et al., 2020; Garcia-Soto et al., 2021). In the last decade, the rate of Arctic Ocean warming was observed to be 4.5 times greater than the global mean (Garcia-Soto et al., 2021). Circulation models indicate a greater increase in average annual surface air temperatures in the Arctic compared to other areas of the planet (Johannessen et al., 2004). This accelerated warming observed in the Arctic is caused by a positive feedback loop fueled by the melting of snow-covered sea ice. As sea ice melts due to increasing atmospheric temperatures, absorption rates of solar radiation increase resulting in even further sea ice melt. This process is called Arctic amplification (Rantanen et al., 2022). It has also been observed that there is a general freshening of the Arctic Ocean and surrounding seas in the last decade due to increases in precipitation, river discharge, and sea ice melt (Chang and Devol, 2009; Peterson et al., 2006). This has the potential to affect deep water formation, sea ice extent and thickness, overall ecosystem function, and the livelihood of surrounding communities (Chang and Devol, 2009; Whitmore et al., 2022). Microbes are often referred to as the first responders in an ecosystem. Short generation times allow microbes to act as markers for environmental changes. They are also likely to be one of the first organisms impacted by very small changes in their local surroundings. Specifically, many microbial niches are situated at the margins between frozen and liquid water, for example, in brine channels in sea ice, permafrost, glacial weathering crusts, or englacial vein boundaries (Edwards et al., 2020). In these micro-environments, liquid water is limited, hindering microbial activity. Thus, even small shifts in temperature can radically alter the niches available to microbial populations (Edwards et al., 2020). Ocean warming may affect these organisms and their productivity in many ways due to changes in sea ice and snow coverage (Box et al., 2019), temperature (Cheng et al., 2017), nutrient availability (Kwiatkowski et al., 2020), competition, pH (Hoegh-Guldberg et al., 2017), and dissolved oxygen concentrations (Nayfach et al., 2020; Schmidtko et al., 2017). As the Arctic continues to change and temperatures rise, marine microbial communities are expected to undergo changes in cell size and structure (MORÁN et al., 2010), biogeography (Falkowski and Oliver, 2007), and food web dynamics (Baer et al., 2014; O’Connor et al., 2009). One group of microbes, called Archaea, are important contributors to biogeochemical cycling but remain relatively uncharacterized in the Arctic Ocean. Using DNA sequencing technologies, Woese and Fox (1977) provided the first evidence that Archaea constituted the third Domain of life alongside Eukarya and Bacteria (Baker et al., 2020; Offre et al., 2013), but more than 40 years later our knowledge of archaeal biology is still limited compared to the other Domains. Archaea are a unique group of single-celled organisms that appear in many ways like Bacteria but have numerous biochemical and genetic features that are more similar to Eukaryotes (Roy Chowdhury and Dutta, 2012). Although it appears that Archaea can be found in all environments, the chemistry of their ecological context is still of great interest. Offre et al. (2013) describe two principal issues in obtaining a full understanding of Archaea: i) identifying the entirety of biogeochemical processes that can be performed by members of the Domain Archaea; and ii) measuring the total archaeal biomass, their productivity, and the nutrient cycling processes performed in their respective environments. Historically, our understanding of microorganisms has been hindered by our inability to cultivate them, and many studies relied on culture-dependent methods (Baker et al., 2020). However, only six of the 27 proposed archaeal phyla currently have cultured representatives (Adam et al., 2017; Baker et al., 2020), making the majority of these organisms identifiable solely by culture-independent molecular studies (Rinke et al., 2013). Microbiomes consist of all the organisms, their genomes, and the surrounding habitat from a defined environment (Frioux et al., 2020). The most common method for culture-independent microbiome analyses utilizes high-throughput DNA sequencing (HTS) with either a targeted approach (amplicon sequencing of a barcode sequence) or an untargeted approach (shotgun metagenomic sequencing) (Frioux et al., 2020; Liu et al., 2020; Teeling and Glockner, 2012). Although more expensive and time-consuming, metagenomic sequencing is more informative compared to amplicon sequencing as it can provide both taxonomic and functional information (Adam et al., 2017; Baker et al., 2020; Liu et al., 2020). Briefly, DNA is sequenced as short reads which are assembled into longer contigs. These contigs are then binned to create partial or complete metagenome-assembled genomes (MAGs). Metagenomic approaches can identify a higher diversity of archaeal lineages than 16S ribosomal RNA gene amplicon sequencing, which relies on commonly used universal primer sets, potentially missing entire clades of organisms (Adam et al., 2017; Liu et al., 2020; Teeling and Glockner, 2012). Metagenomic binning is a relatively new development in the world of bioinformatics and is typically the sole source of genetic information for uncultured organisms (Frioux et al., 2020). A considerable hurdle for metagenomic studies is determining the origin of the derived genome bins (Rinke et al., 2013), especially for assembling low-abundance and closely related members of the community (Bickhart et al., 2022). This often depends on the sequencing depth and quality. It becomes very difficult to assemble genomes closely related to a highly abundant species as those bins are often combined. It is possible to obtain high-quality MAGs but typically only after a large amount of manual polishing and curation. If a community has many members, the problem of creating reliable MAGs quickly becomes an overwhelming task. Binning algorithms have undergone many advancements that include better clustering approaches of contigs and their abundance profile, GC content, and k-mer content (Frioux et al., 2020). GC content refers to the proportion of guanine (G) and cytosine (C) in the genomes of the microbes. In bioinformatics, k-mers refer to substrings of length k contained within a biological sequence, for example, a k-mer of length four (4-mer) would represent a DNA sequence that is four bases long (e.g., ATTC). The ability to reconstruct complete or nearly complete MAGs is influenced by numerous factors such as microbial diversity, biomass, sequencing effort, read length, intrapopulation sequence, gene diversity, and the assembly and binning algorithms used (Meziti et al., 2021). Reconstructing these genomes becomes extremely important when trying to understand the small-scale changes that are occurring in a specific environment (Frioux et al., 2020). The majority of uncultured genomes originate from sediments and water columns and represent a sizeable proportion of microbial life (Adam et al., 2017; Karner et al., 2001; Offre et al., 2013; Reji et al., 2019) making the Arctic Ocean a desirable location for researching microorganisms. Developing a unifying concept of life is often hindered by a lack of available data, especially from microbes in cold environments. Understanding the genomic data and metabolic potential of archaeal representatives in the Arctic Ocean is an important step toward understanding their role in the environment and providing a new perspective on their metabolic diversity. Two primary archaeal phyla, the Euryarchaeota and Crenarchaeota, were initially proposed based on phenotypic characteristics (Baker et al., 2020; Offre et al., 2013), but analysis of genomic data and advances in sequencing technologies, including HTS and metagenomic assembly and binning, have had a strong impact on archaeal systematics leading to the description of 20 additional phyla and multiple new clades in the last decade (Adam et al., 2017; Baker et al., 2020). Currently, there are three superphyla recognized as the Asgard, DPANN, and TACK (Baker et al., 2020; Guy and Ettema, 2011; Rinke et al., 2013). The DPANN and TACK superphyla were originally named based on the phyla present within them: DPANN referring to the Diapherotrites, Parvarchaeota, Aenigmarchaeota, Nanohaloarchaeota, and Nanoarchaeota; and TACK to the Thaumarchaeota, Aigarchaeota, Crenarchaeota, and Korarchaeota. The TACK Superphylum is one of the most extensively studied clades today (Adam et al., 2017; Baker et al., 2020). Thaumarchaeota is thought to be one of the most numerically abundant clades of Archaea on the planet, nearing 20% of total microbes in some deep ocean and soil environments (Baker et al., 2020; Brochier-Armanet et al., 2008; Karner et al., 2001; Reji and Francis, 2020; Reji et al., 2019; Santoro et al., 2011; Spang et al., 2015). A highly studied representative of this group, *Nitrosopumilus maritimus*, has adapted to life under extreme nutrient limitations and high specific oxidation rates of ammonium at concentrations found in open oceans (Martens-Habbena et al., 2009). Currently, one phylum, the Euryarchaeota, does not fall into a superphylum but contains the greatest diversity of cultured lineages including members of the classes Methanopyri, Methanococci, Methanobacteria, Methanomicrobia, Archaeoglobi, Halobacteria, Thermococci, and Thermoplasmata (Baker et al., 2020; Offre et al., 2013). Euryarchaeota are broadly associated with diverse metabolisms including methanogenesis, methane-oxidation, denitrification, sulfate reduction, iron oxidation, and organotrophy (Offre et al., 2013). Culture-independent studies have shown that these organisms are important contributors to global biogeochemical cycling, such as the production of methane and the remineralization of nitrate (Baker et al., 2020; Offre et al., 2013). However, these classifications are always evolving. Rinke et al. (2020) proposed a major reclassification of the Domain consisting of 16 phyla, combining the TACK superphylum into one, and separating the Euryarchaeota into five new phyla. The first environmental archaeal sequences detected in marine environments were separated into two groups: Marine Group I (MG-I), now known as Thaumarchaeota, and Marine Group II (MG-II), within Euryarchaeota (Brochier-Armanet et al., 2008; Yilmaz et al., 2016). This structure has been further expanded to contain three main marine groups. Within Euryarchaeota, two groups were identified: Poseidoniales or MG-II (Pereira et al., 2019; Rinke et al., 2020), and Pontarchaea or Marine Group III (MG-III) (Baker et al., 2020; Yilmaz et al., 2016). Currently, MG-II representatives are not well understood as there are no cultured representatives from this group (Pereira et al., 2019). 16S rRNA gene sequences have revealed that MG-II may be divided into two main monophyletic groups, the MG-IIa, now known as *Candidatus* Poseidonaceae, and the MG-IIb, now known as *Candidatus* Thalassarchaeaceae (Rinke et al., 2020). The MG-III are commonly found in oceans worldwide, however, they are typically found in very low abundances. They have been shown to contain genes for recycling high molecular weight organic compounds (Baker et al., 2020), but the majority of their metabolic functions are unknown. For the remainder of the text, the archaeal clades of interest will be referred to by their marine group name (MG-I; MG-II; MG-III). Certain groups of Archaea have been found to preferentially reside in colder environments, often lower in the water column (Pereira et al., 2019), while others have shown a preference for warmer photic zones (Rinke et al., 2019). MG-I are highly abundant in deeper waters below the thermocline (Ionescu et al., 2009), while MG-II often surpass MG-I abundances in surface waters, especially in summer months (Pereira et al., 2019; Yilmaz et al., 2016). The MG-IIa (*Candidatus* Poseidoniaceae) is typically composed of surface microbes, while the MG-IIb (*Candidatus* Thalassarchaeaceae) are often found at depths below 200 m (Pereira et al., 2019). Most of these studies did not sample below 500 m. These groups have been shown to have different seasonal variations depending on their location of origin (Pereira et al., 2019). For example, MG-IIa are found in higher abundance in the polar oceans in winter but were predominant in summer months in the Mediterranean (Pereira et al., 2019). It has been proposed that MG-IIa have evolved to occupy many diverse niches while remaining surface residing (Rinke et al., 2019). Some rarer phyla have also been identified in deep-sea waters including Bathyarchaeota and Woesearchaeota from the Indian Ocean and North Pacific Ocean (Pereira et al., 2019). Liu et al. (2018) collected samples off the coast of Georgia, USA from April to November 2014. Nitrite was found at low concentrations in the surface waters but increased below *∼*100 m (Liu et al., 2018). Consistent with other studies (Ionescu et al., 2009; Pereira et al., 2019; Reji et al., 2019; Yilmaz et al., 2016), the majority (80%) of MG-I 16S ribosomal RNA gene sequences were recovered at 200 m (Liu et al., 2018). These results contrast with studies from high-latitude regions such as the North Sea (Herfort et al., 2007; Pitcher et al., 2011; Wuchter et al., 2006) and polar oceans (Church et al., 2003) where MG-I abundance was highest at even greater depths (>200 m). Liu et al. (2018) proposed that these differences in abundances may be attributed to the enhanced vertical mixing that occurs at lower latitudes in summer months. Although the MG-I abundance increased around 200 m, Euryarchaeota remained the dominant archaeal phylum in all the samples (Liu et al., 2018). This is consistent with results from the northeastern South China Sea (Liu et al., 2017). It is commonly suggested that the distribution of MG-I may be related to photoinhibition, as light may be an inhibitor for growth, however, it is not clear if this inhibition is a result of light directly (phototaxis) or if photochemical reactive oxygen species play a role (Merbt et al., 2011; Qin et al., 2014). This may also explain the decrease of MG-I in the summer months in the polar zones, as the levels of sunlight increase dramatically during that time of year, and their preference is to reside deeper within the water column. Liu et al. (2018) also suggest that at sites where light penetration was consistently low, the water temperature was a secondary control on the bloom potential. Seasonal variability in the Arctic Ocean will likely significantly impact the communities of microbes residing there: this ocean is ice-covered during much of the winter and complete darkness can last up to two months (Christman et al., 2011). These conditions are expected to support the growth of ammonia-oxidizing Archaea (AOA) as shown by other studies (Baer et al., 2014; Christman et al., 2011; Liu et al., 2018). Arctic stratification due to salinity is a key factor influencing circulation in the Arctic Ocean, ice coverage, and productivity (Farmer et al., 2021). Surface waters tend to be nutrient-poor and colder than their underlying warmer, nutrient-rich water (Farmer et al., 2021). The PML tends to be comprised of colder, fresher water where primary production typically takes place (Oziel et al., 2022). The subsurface is comprised of the saltier, nutrient-replete waters of Pacific and Atlantic origin (Farmer et al., 2021; Oziel et al., 2022). Water inputs into the Arctic Ocean are supplied from distinct Atlantic and Pacific sources. Salty Atlantic waters originate from inflows through the Fram Strait and the Barents Sea. Atlantic waters dominate the eastern and central Arctic Ocean. In the Western Arctic, this Atlantic water is covered by fresher Pacific water coming in from the Bering Strait (Farmer et al., 2021). Stratification is stronger in the Western Arctic as compared to the eastern Arctic (Farmer et al., 2021). The PML stratification is due to excess precipitation, river outflow/discharge, sea ice melt, and the advection of low-salinity ocean waters across the Bering Strait (Farmer et al., 2021). Understanding the various water masses and the distribution of different microbial clades can provide insight into the evolution and ecology of the organisms as they travel through the Arctic Ocean. Environmental variables such as light, nutrient availability, water column mixing, and water mass characteristics may control the presence or absence of different microbial clades (Arandia-Gorostidi et al., 2022). Genome-resolved metagenomics offers an opportunity to identify the causative factors for these trends at the genomic level, for example, due to the presence or absence of particular genes and pathways, or proteomic adaptation to low temperature. To investigate the ecology of Archaea in the Western Arctic Ocean, metagenomic seawater samples were collected during the GEOTRACES 2015 expedition. Samples were collected in late summer from 66 stations in the Western Arctic Ocean (Figure 1). Physical parameters collected included depth, temperature, salinity, nutrient concentrations, and heavy metal concentrations. After sample filtration, DNA extraction, and DNA purification, metagenomic sequencing was performed. The goals were to catalogue and identify the archaeal taxa collected from the Western Arctic and to identify the population structure of archaeal communities in the Western Arctic using metagenome-assembled genomes (MAGs). Previous studies have suggested that temperature and light penetration are the main factors driving archaeal abundance patterns, however, we hypothesized that water masses in the Arctic Ocean would be the strongest factor influencing archaeal community structure. Each water mass layer has different physical and chemical properties that are likely to influence any abundance variations observed. For functional convenience, however, the layers themselves will be used as a proxy for the underlying chemical and physical properties that are driving the abundance distributions observed. Based on previously collected 16S rRNA gene sequence data from the Chukchi Sea (Supplementary Figure S1), we expected to recover MAGs of commonly occurring taxa, including *Nitrosopumilus* and other members of MG-I, as well as members of MG-II and MG-III.

**Figure 1:**
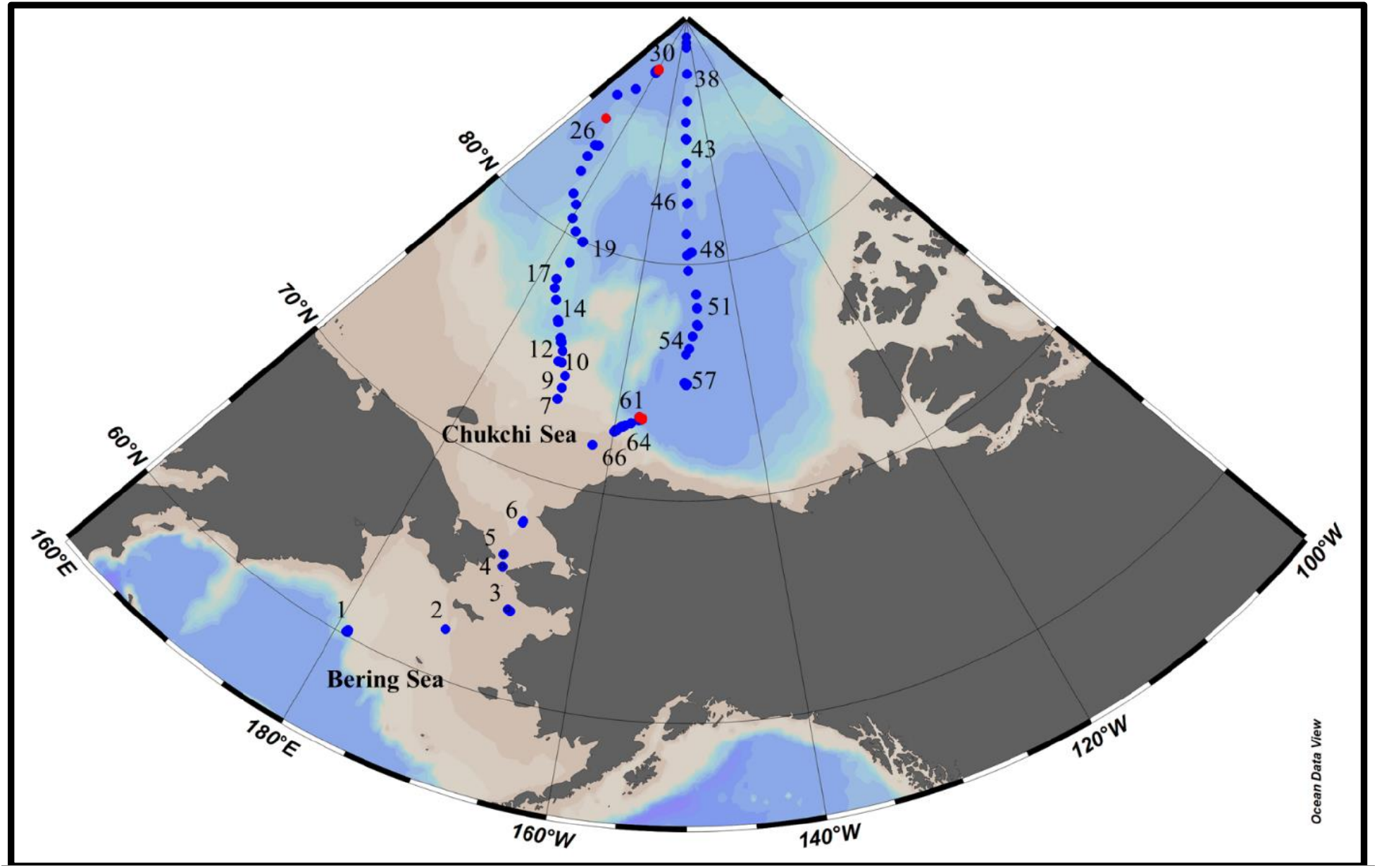
Geographical map displaying the stations from the 2015 GEOTRACES expedition. Red indicates the stations (26, 30, 60, 61) where metagenomic data was collected.

**Table 1:** Classification of the archaeal groups referred to in the text.

| Superphylum/phylum | Clade | Marine Group |
|---|---|---|
| TACK | Thaumarchaeota | I |
|  | Aigarchaeota | – |
|  | Crenarchaeota | – |
|  | Korarchaeota | – |
| DPANN | Diapherotrites | – |
|  | Parvarchaeota | – |
|  | Aenigmarchaeota | – |
|  | Nanohaloarchaeota | – |
|  | Nanoarchaeota | – |
| Asgard | – | – |
| Euryarchaeota | Poseidoniales | II |
|  | Pontarchaea | III |

## 2 Materials and Methods

### 2.1 Sample collection and environmental data

An oceanographic expedition (HLY1502; Kadko and Landing, 2015) that departed from Dutch Harbor, Alaska, was conducted from August to October in the Western Arctic as part of the GEOTRACES program (GN01) which mainly focused on characterizing the distribution of trace elements and other chemical parameters in the oceans. GN01 transited through the Bering and Chukchi Seas to the North Pole in two transects (Figure 1): 1) through the Makarov Basin; and 2) through the Canada Basin. Microbial community seawater samples were collected in the Western Arctic Ocean during this expedition using a CTD rosette. The collected water was filtered onto 0.22 µm Sterivex filters using a peristaltic pump, allowing sample water to run through the tubing equipment before filter attachment. As much water as possible was removed from the filters using a syringe and then stored at *−*80 °C until sequencing could be performed. At each sampling site, depth, salinity, and temperature were measured using a CTD cast. Other variables including particulate organic nitrogen (PON), particulate organic carbon (POC), nitrate (NO_3_^−^), nitrite (NO_2_^−^), ammonium (NO_4_^+^), and dissolved oxygen (DO) were also measured according to the GEOTRACES protocols (Cutter et al., 2017). Briefly, water was collected via the ship’s rosette using approximately 250 mL per depth (50 mL in triplicate, plus bottle rinses). Samples were filtered via a pressure filtration using a 0.22 µm Sterivex filter into 60 mL square wide-mouth HDPE bottles and stored at *−*20 °C. Dissolved oxygen was measured using the automated Winkler method with a detection limit of 1 µmol L^−1^. Ammonia, nitrite, and nitrate were measured using an automated colorimetric method with detection levels of 0.1 µmol L^−1^. Silicate was measured in the same way with a detection level of 0.4 µmol L^−1^.

### 2.2 Metagenomic sequencing, assembly, and binning

DNA was extracted and purified from filters using the Qiagen PowerWater Sterivex kit. DNA concentration was measured using a Qubit and sequenced on an Illumina MiSeq at 2*×*250 base pairs (bp). The following metagenomic pipeline is shown in Figure 2. BBtools (Bushnell et al., 2017) was used to remove unwanted DNA sequences (i.e., barcodes, adaptors, sequencing artifacts, and low-complexity sequences) and perform quality controls using the following commands: optical duplication removal (clumpify.sh); low-quality region removal (filterbytile.sh); adapter trimming, and sequencing artifact and low-entropy sequencing removal (bbduk.sh); error correction (bbmerge.sh, clumpify.sh, tadpole.sh); and normalization (bbnorm.sh). Reads were additionally normalized (bbnorm.sh) before co-assembly. Clustering with minhash was performed for the co-assembly. A co-assembly with 128 samples was performed using metaSPAdes (Nurk et al., 2017) to elucidate the lower abundance members within each sample. Error-corrected reads were mapped back to the co-assembly (bbmap.sh; percent identity = 100%) to obtain the relative abundances of each contig in each sample. Metagenomic binning was performed by placing contigs into putative microbial genomes (metagenome-assembled genomes or MAGs) using MetaBAT2 (Kang et al., 2019), MaxBin2 (Wu et al., 2015), and MetaBinner (Wang et al., 2023). The binning outputs were consolidated using DAS Tool (Sieber et al., 2018), which is an automated method that integrates the results of numerous binning algorithms to calculate an optimized, non-redundant set of bins from a single assembly. Completeness and contamination measures were determined using CheckM (Parks et al., 2015). To do this, CheckM uses Prodigal (Hyatt et al., 2010) to identify genes in a sequence by first placing the genome on a tree and using that phylogenetic placement to identify sets of genes that should be in the genome. It then looks for those genes within the genome using HMMER, which uses profile hidden Markov models to identify sequence homologs. The completeness is an estimate of the fraction of genes that are expected to be there which are actually found. The contamination estimate is based on identifying the number of single copy genes that should only be there once.

**Figure 2:**
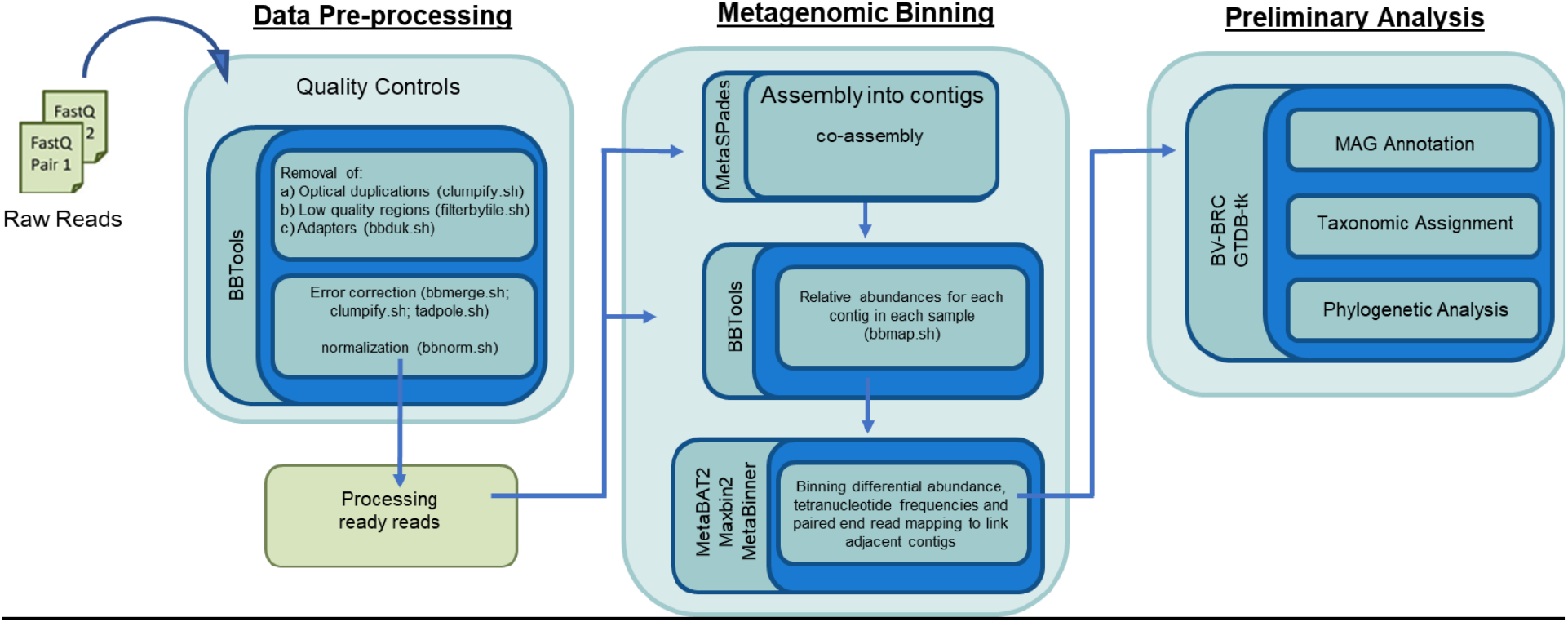
Metagenomic pipeline used in the analysis.

### 2.3 MAG annotation and taxonomic assignment

MAGs were annotated using the whole genome analysis tool available from Bacterial and Viral Bioinformatics Resource Center (BV-BRC), formerly called PATRIC, which is an online portal for genomes and metagenomics (Davis et al., 2019). BV-BRC uses the RAST toolkit (RASTtk) (Brettin et al., 2015) to identify open reading frames, translate open reading frames into amino acid sequences, compare amino acid sequences to databases of known proteins, and assign functional categories to each gene in the genomes. BV-BRC was also used to calculate other bin quality measurements like the N50 and L50. Contig N50 is the sequence length of the shortest contig at 50% of the total assembly length and captures how much of the genome is covered by large contigs. Contig L50 is the number of contigs that are equal to or longer than the contig N50 value. Therefore, bins with a high N50 and low L50 are considered to be of higher quality. Bins were tentatively classified using Minhash Sketching (sendsketch.sh) and the similar genome finder tool in BV-BRC. Taxonomic assignment was completed using the Genome Taxonomy Database Toolkit (GTDB-Tk; Chaumeil et al., 2022). GTDB-Tk is a computational toolkit that provides taxonomic classification of bacterial and archaeal genomes by placing them into a domain-specific, concatenated protein reference tree. Briefly, genomes are assigned to a domain (Bacteria or Archaea) with the highest proportion of shared marker genes. The marker genes are then aligned using HMMER and concatenated into a single multiple sequence alignment (MSA) which is then placed into a domain-specific reference tree using pplacer (Matsen et al., 2010). The taxonomic classification of a query genome is based on its placement in the reference tree, as well as its Average Nucleotide Identity (ANI) to reference genomes. Phylogenetic analysis was performed using the phylogenetic tree tool in BV-BRC (50 conserved single-copy core genes, allowing for up to 10 duplicates and 10 deletions). The phylogenetic tree tool uses the codon tree method which analyzes aligned protein and coding DNA sequences of single copy genes using protein global families as homology groups using the RAxML program (Stamatakis, 2006). Trees were rooted and manually curated using iTOL (Letunic and Bork, 2006), which is an online tool to display, annotate, and manage phylogenetic trees.

### 2.4 Water mass identification and biogeographic mapping

Ocean Data View (Schlitzer, 2023) was used in the creation of section plots for various parameters (temperature, salinity, dissolved oxygen concentrations, etc.) using the default DIVA gridding settings. Contours were added and colour mapping was auto adjusted. Pacific Water identification was performed using measured silicate concentrations and a linear combination of nitrate and phosphate concentrations (*N^*^* = 0.87 [*N −* 16*P* + 2.9]; Gruber and Sarmiento, 1997). The latter equation is used to measure the distribution of nitrification and denitrification, where N denotes nitrate concentrations (µmol kg^−1^) and P denotes phosphate concentrations (µmol kg^−1^). In this equation, the constant (2.9 µmol kg^−1^) was determined by forcing the global mean of N* based on the Geochemical Ocean Sections Study (GEOSECS) data (1972-1978) to be zero. This equals the value that would be calculated using the GEOSECS-based global mean nitrate of 30.38 µmol kg^−1^ and global mean phosphate of 2.08 µmol kg^−1^. This equation uses values of stoichiometric N:P ratios for the mineralization of organic matter (the Redfield ratio of 16:1; Redfield et al., 1963). This ratio is derived from the global average of biomass in the oceans due to primary and secondary productivity. Lower values indicate nitrate depletion relative to Redfield ratios due to microbial denitrification, in which nitrate is reduced via nitrite to the gases nitric oxide (NO), nitrous oxide (N_2_O), and dinitrogen (N_2_), which can be lost to the atmosphere. Therefore, a lower N* value represents a state where the nitrate depletion is greater relative to what would be expected if nitrate was only being utilized for the production of biomass. The depth of the PML was identified by locating the temperature minimums of each cast. Temperature and salinity measurements were used in the determination of all other water masses following Pisareva et al. (2015). R (R Core Team, 2021) in RStudio was used to map the distribution of taxa in relation to environmental parameters such as depth, temperature, salinity, light, and nutrients. Relative abundance was used in this analysis, which is a measure of the proportion of total microbial abundance and does not capture the absolute cell counts or abundances. Therefore, an increase in relative abundance could be caused by a decrease in the abundance of other organisms while the absolute counts may remain the same throughout the water column. Statistical analyses were done in R using the vegan package (Oksanen et al., 2022).

## 3 Results

### 3.1 Water mass determination and physical parameters

The GEOTRACES 2015 Expedition (GN01) took place in late summer departing from Alaska. In total 66 stations were measured for temperature, salinity, depth, and nutrient concentrations. Of the 66 stations, the deepest were stations 30 (3951 m), 32 (4239 m), 34 (4221 m), and 56 (4394 m), which were closest to the North Pole. The shallowest sampling depths were from stations 3 (33 m), 2 (49 m), and 66 (34 m) along with other stations from the southern coastal regions and the Chukchi Sea. Stratification due to temperature and salinity was observed in the northern regions, however, it was more difficult to detect stratification in the southern coastal regions of the Bering Strait and Chukchi Sea using the same metrics. A linear combination of nitrate and phosphate concentrations (Gruber and Sarmiento, 1997) was used to measure the distribution of nitrification and denitrification, as described in Section 2.4. N* measurements were used to identify the Pacific Ocean water coming through the Bering Strait and over the shelf (Figure 3). Negative N* values are indicative of a sink of N* due to benthic denitrification and indicate Pacific Ocean water coming into the Western Arctic Ocean through the Bering Strait and Chukchi Sea (Deutsch et al., 2001; Gruber and Sarmiento, 1997) and can be seen between approximately 25 m and 150 m up to 84 °N. In the southern coastal areas (<75 °N), there was an increase in the N* values observed in the uppermost 25 m. This tool for determining water masses is not effective in surface waters due to the added influence of other microbial processes that interfere with nutrient content. Therefore, in the surface waters, temperature was used as the main determining factor of water mass differentiation. Another common tracer of Pacific Ocean water entering the Arctic Ocean is elevated silicate concentrations (Jensen et al., 2019; Jones and Anderson, 1986). Silicate concentrations (Figure 4) were observed to be elevated in the same region as N* values were lower. It was observed that the low silicate surface layer corresponded with higher N* values, increased temperature, and decreased salinity in the upper 25–50 m. The temperatures in this region were much warmer, indicating the summer surface layer (SSL). Temperatures, which ranged from *−*1 to 14 °C, were warmest in the southern regions (Figure 5) and displayed a thermocline in the northern regions (>75 °N). The temperature minimums for each cast were used to determine the bottom of the Polar Mixed Layer (PML). The PML near the North Pole was colder than its underlying water masses (*∼−*1 °C). Since the PML is often derived from Pacific water moving into the Arctic in the study region, they will be classified together as the PML in this analysis. Salinity ranged from 25.01 PSU – 35.96 PSU throughout the region (Figure 6). This designation is appropriate as the PML salinities observed here were similar to Jensen et al. (2019). The water below the PML consisted of the more saline Atlantic water and was observed approximately between 200 m and 1000 m. For this analysis, both the Fram Strait Branch (FSB) and the Barents Sea Branch (BSB) have been grouped together. The water below the BSB was characterized as the Arctic Ocean Basin water. Using temperature, salinity, silicate, and N* measurements the water masses observed were classified as SSL, PML, Atlantic water (AW), and Arctic Ocean basin water (ABW) (Figure 7). Dissolved oxygen concentrations ranged from 167.11 to 572.40 µmol kg^−1^ and were lower in the southern coastal regions, likely caused by increased heterotrophic microbial activity, as well as in the ABW (Figure 8). The upper 50 m of the higher latitude stations had higher concentrations of dissolved oxygen, suggestive of oxygen production by photosynthesis and uptake from the atmosphere. Nitrate ranged from 0 (below detection) to 17.6 µmol L^−1^ and nitrite ranged from 0 (below detection) to 1.03 µmol L^−1^. Nitrate and nitrite concentrations were lower in the surface waters for stations below 75 °N and increased below 20 m (Figures 9 and 10). Nitrite concentrations were lower than nitrate at all stations and were essentially zero in the ABW. Ammonia concentrations ranged from 0.002 to 8.700 µmol kg^−1^. Phosphate ranged from 0.168 to 2.816 µmol kg^−1^. Silicate ranged from 0.6 to 61.2 µmol kg^−1^.

**Figure 3:**
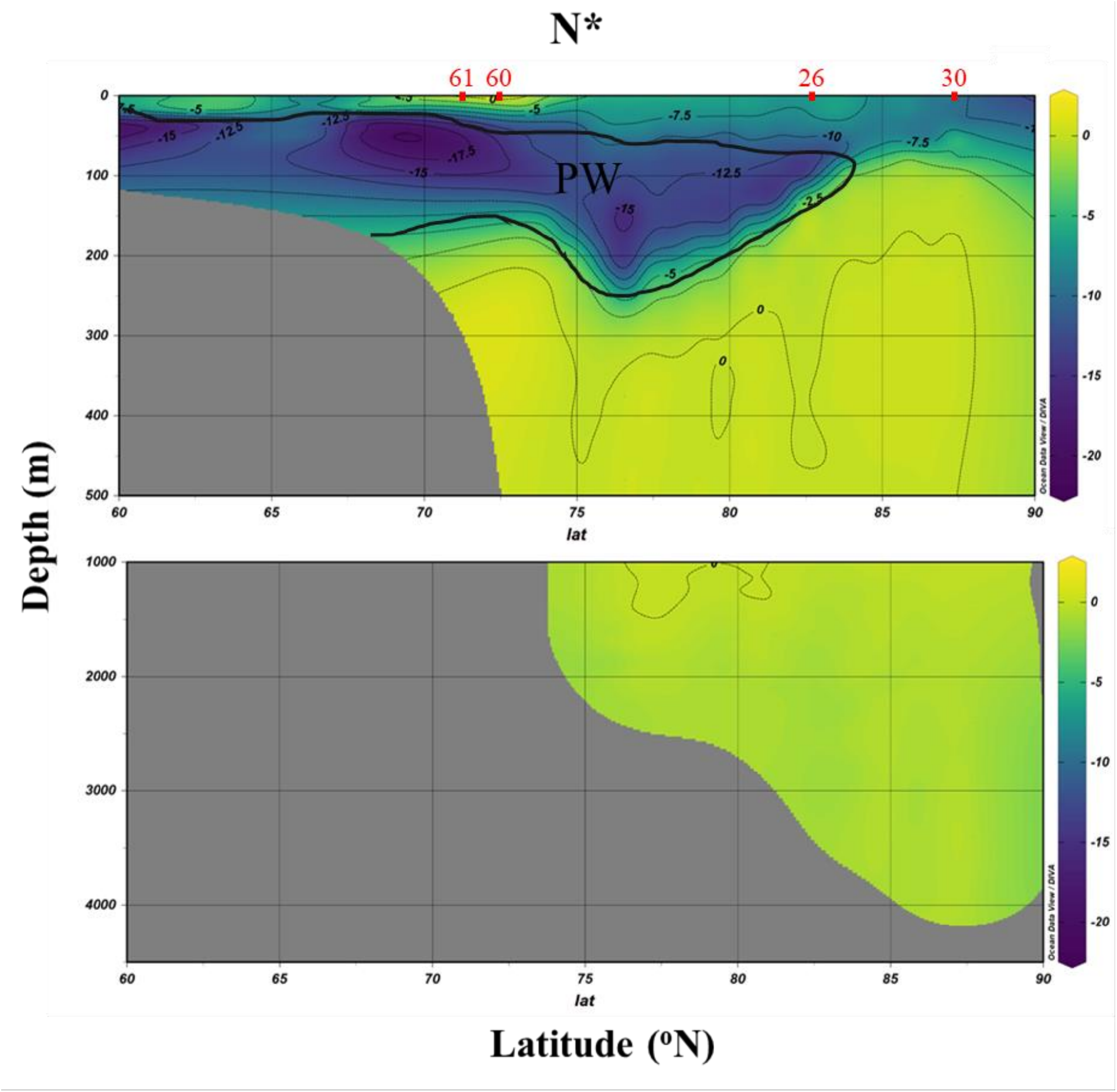
Section plot of N* values for the Western Arctic Ocean during the GEOTRACES 2015 expedition (GN01). Pacific Ocean water is indicated in dark blue as N* values are expected to be lower. The inferred Pacific Ocean water layer is highlighted in black. Values were gridded using weighted-average gridding in ODV. Metagenomic sampling data was collected at stations listed at the top (red).

**Figure 4:**
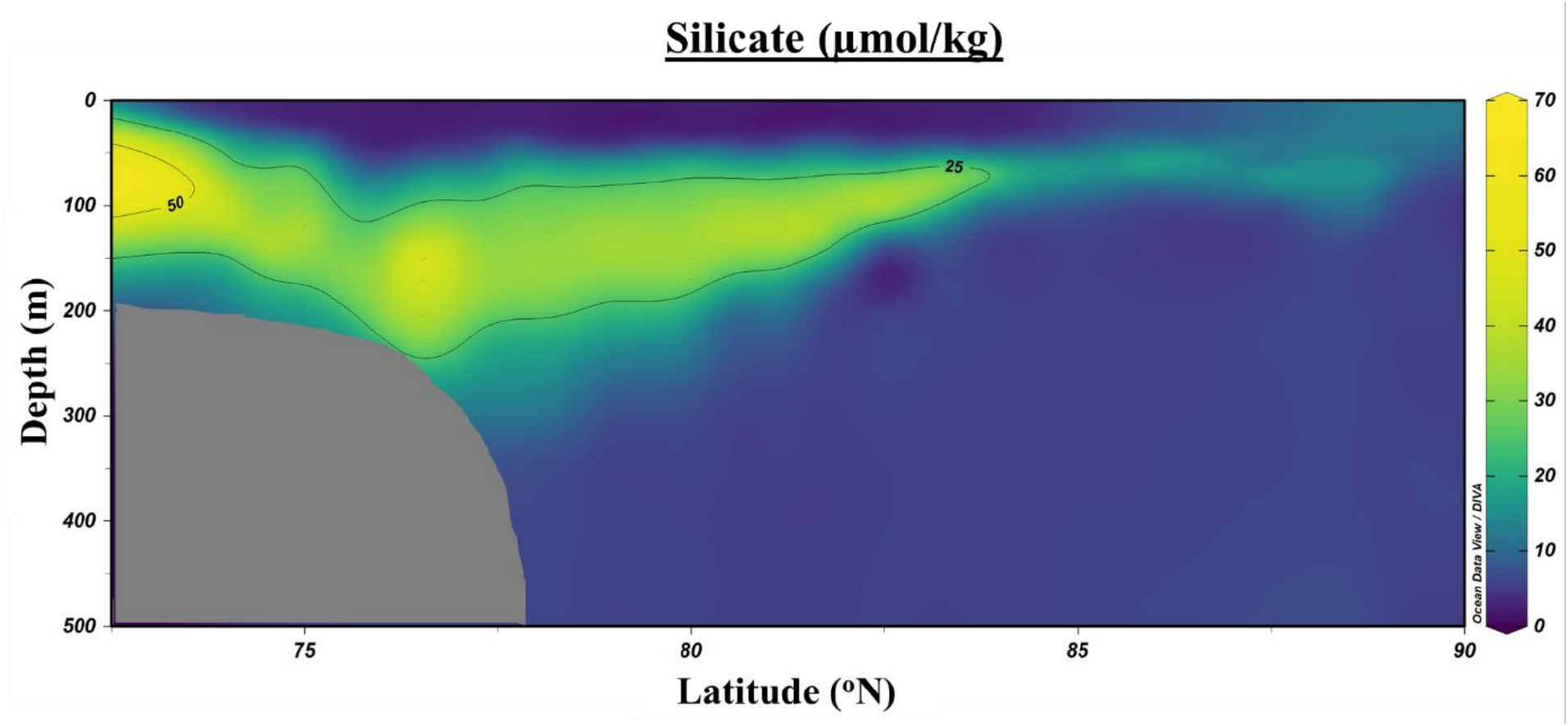
Section plot of silicate measurements for the upper 500 m of the Western Arctic Ocean during the GEOTRACES 2015 expedition (GN01). Values were gridded using weighted-average gridding in ODV.

**Figure 5:**
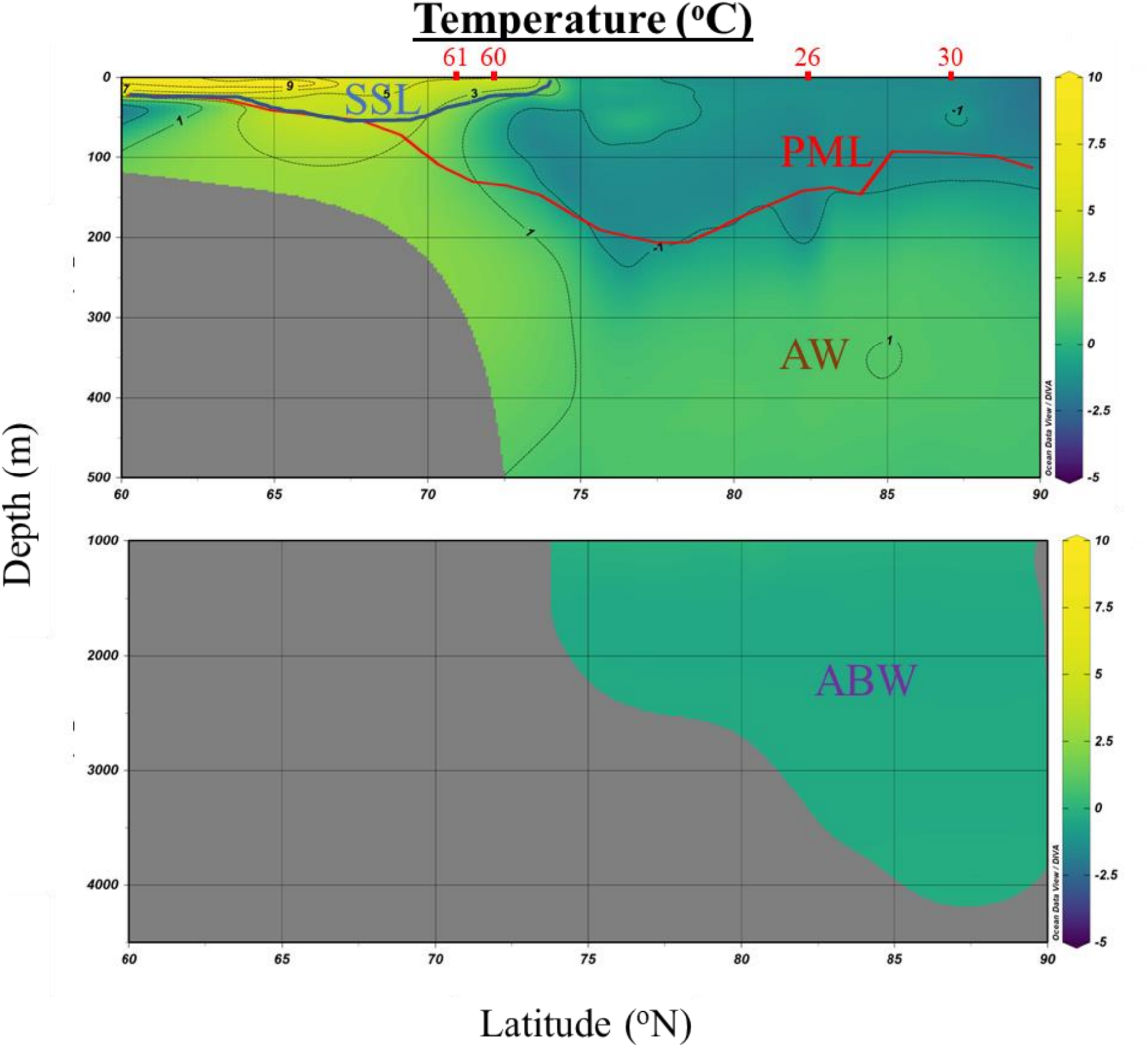
Section plot of temperature measurements for the Western Arctic Ocean during the GEOTRACES 2015 expedition (GN01). Temperature minimums for each cast were used to determine the bottom of the Polar Mixed Layer (PML; red). The summer surface layer (SSL; light blue) is derived from warmer temperatures in the south. The Atlantic water (AW; brown) is warmer than the PML above and warmer than the Arctic Ocean Basin water (ABW; purple) layer below. Values were gridded using weighted-average gridding in ODV. Metagenomic sampling data was collected at stations listed at the top (red).

**Figure 6:**
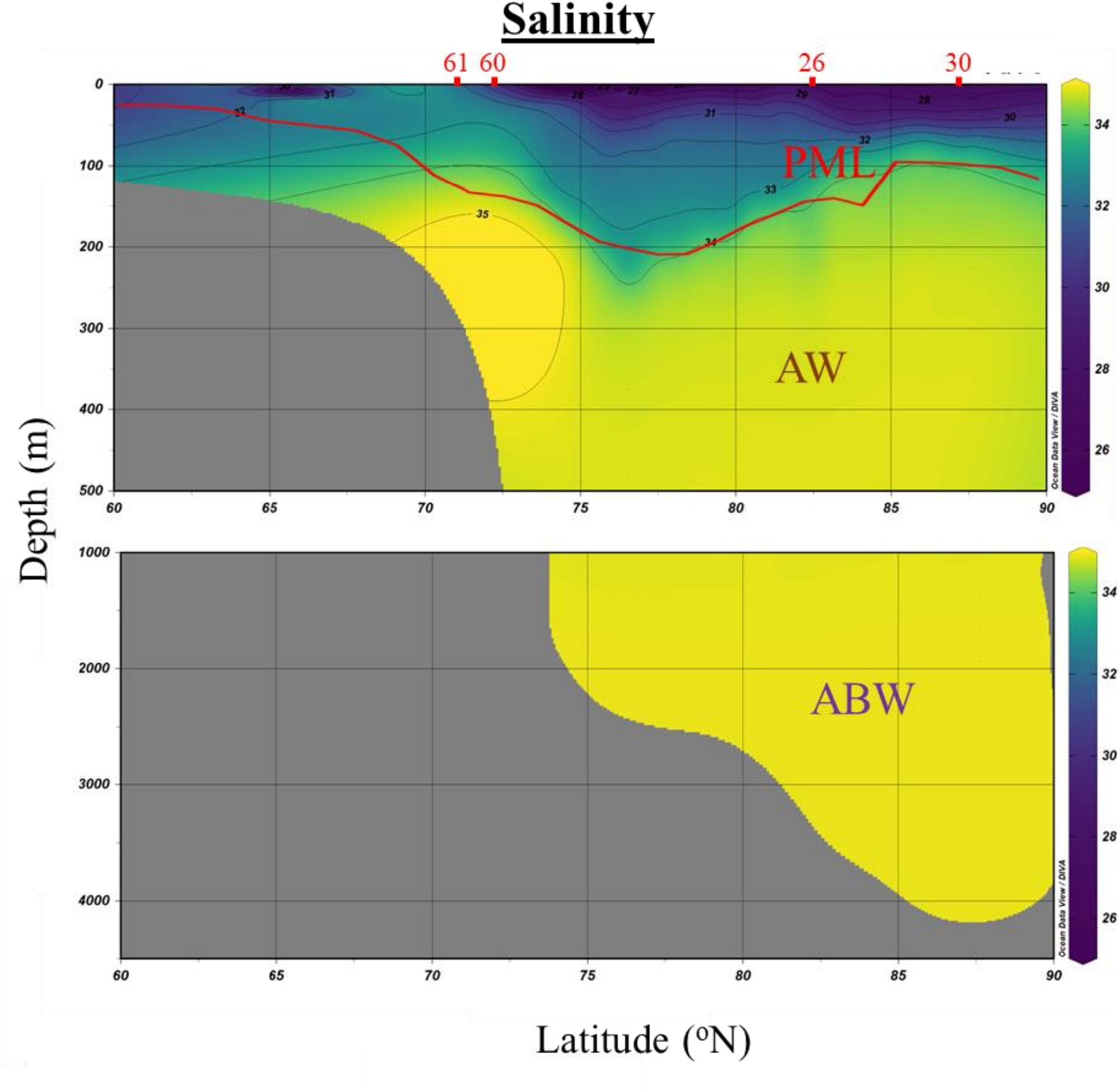
Section plot of salinity measurements for the Western Arctic Ocean during the GEOTRACES 2015 expedition (GN01). Temperature minimums for each cast were used to determine the bottom of the Polar Mixed Layer (PML; red). The Atlantic water (AW; brown) and Arctic Ocean Basin water (ABW; purple) are much more saline than the PML above. Values were gridded using weighted-average gridding in ODV. Metagenomic sampling data was collected at stations listed at the top (red).

**Figure 7:**
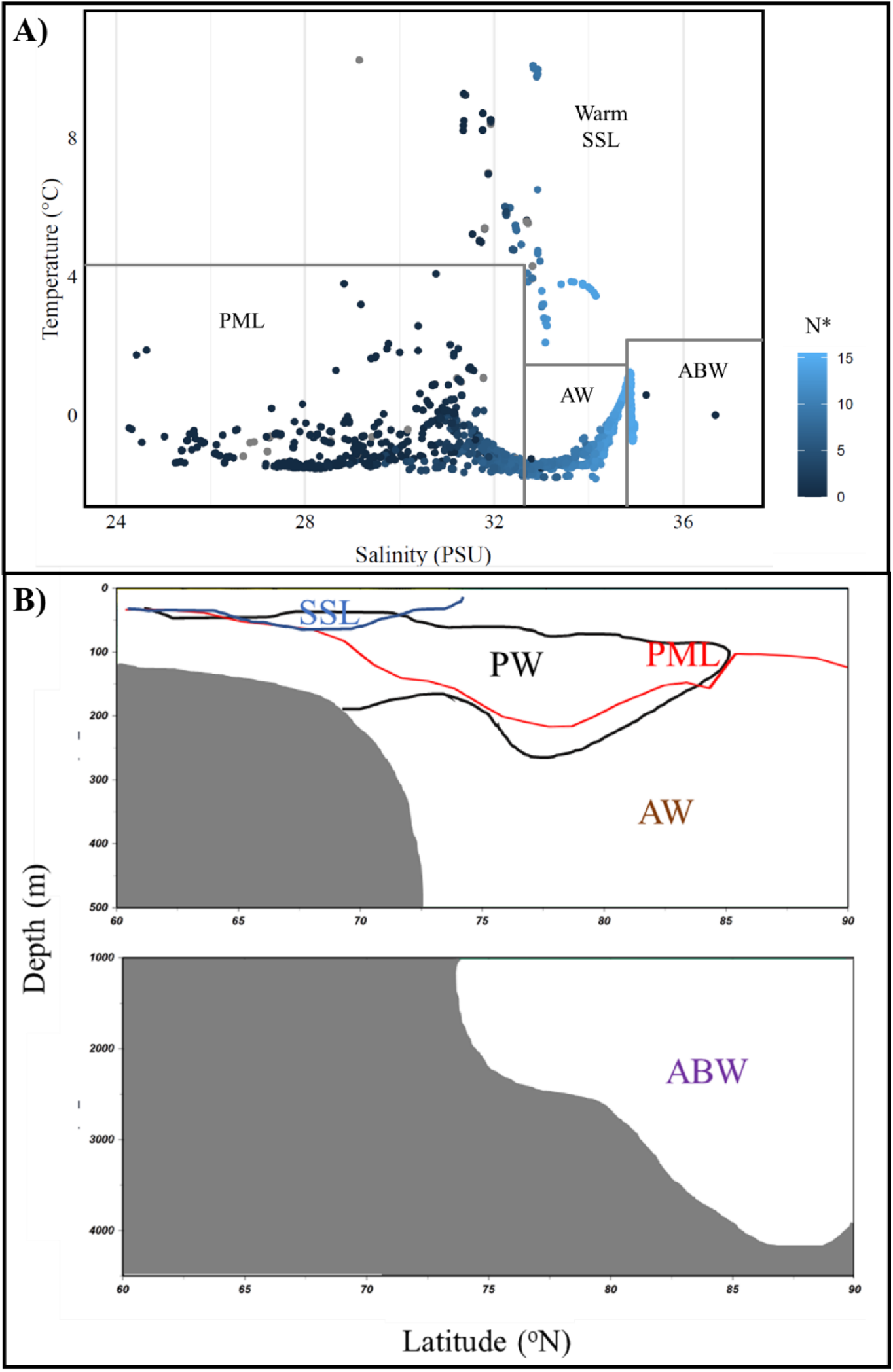
Water mass determinations: A) Temperature-Salinity plots of water sampled; B) Section plot of estimated water layers during the GEOTRACES 2015 expedition (GN01) designating the water masses observed.

**Figure 8:**
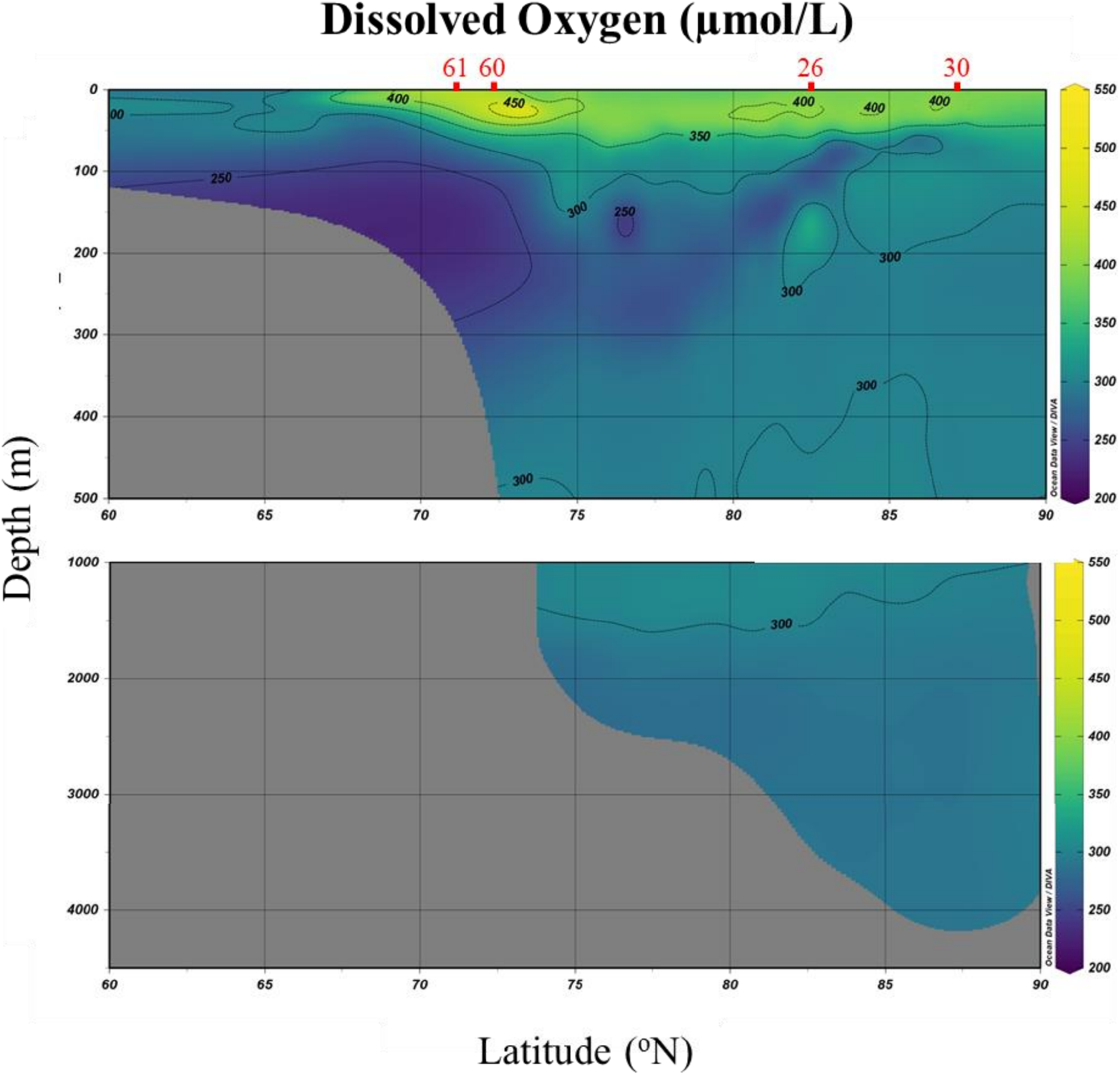
Section plot of dissolved oxygen concentrations for the Western Arctic Ocean during the GEOTRACES 2015 expedition (GN01). Values were gridded using weighted-average gridding in ODV. Metagenomic sampling data was collected at stations listed at the top (red).

**Figure 9:**
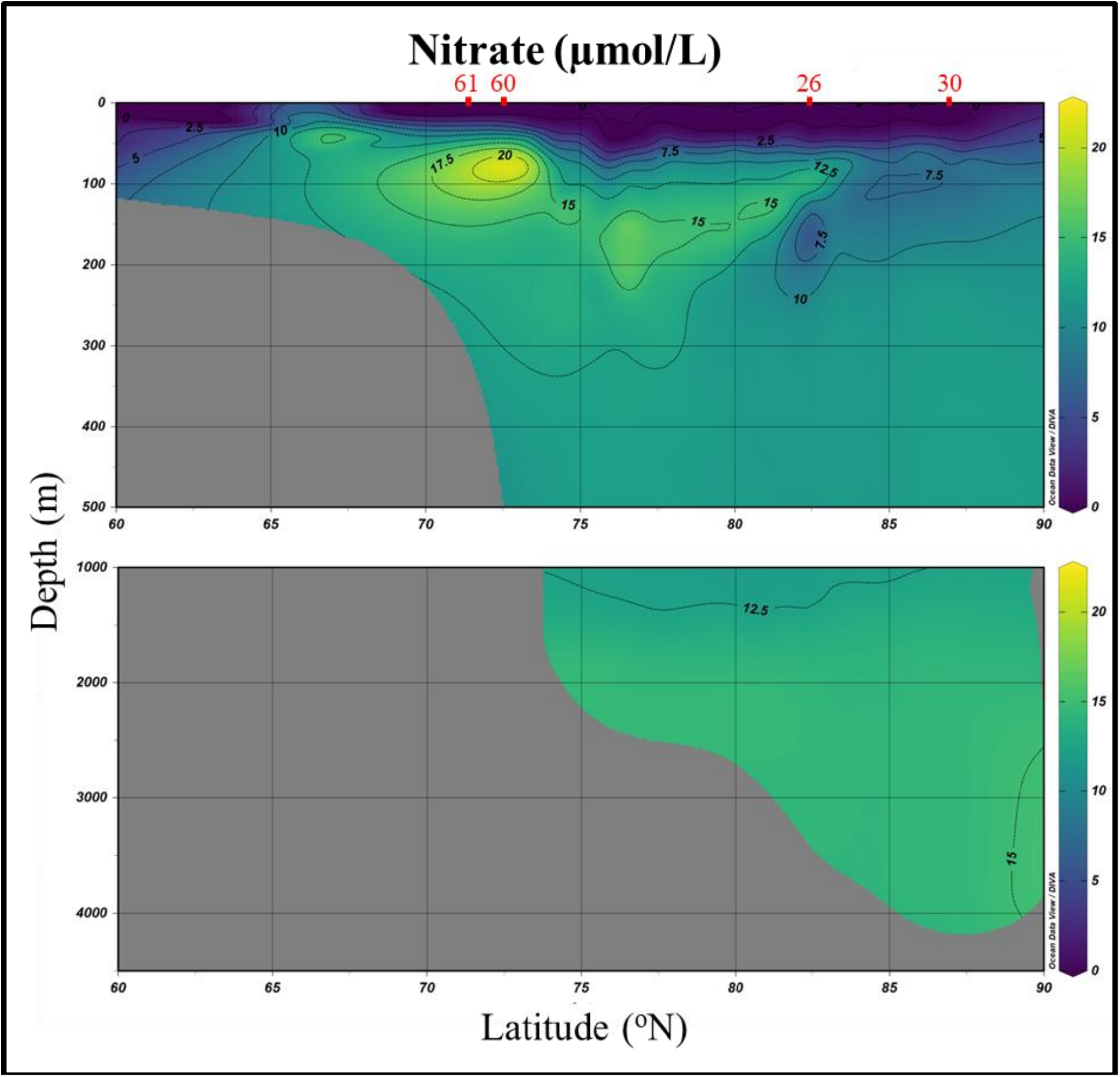
Section plot of nitrate concentrations for the Western Arctic Ocean during the GEOTRACES 2015 expedition (GN01). Values were gridded using weighted-average gridding in ODV. Metagenomic sampling data was collected at stations listed at the top (red).

**Figure 10:**
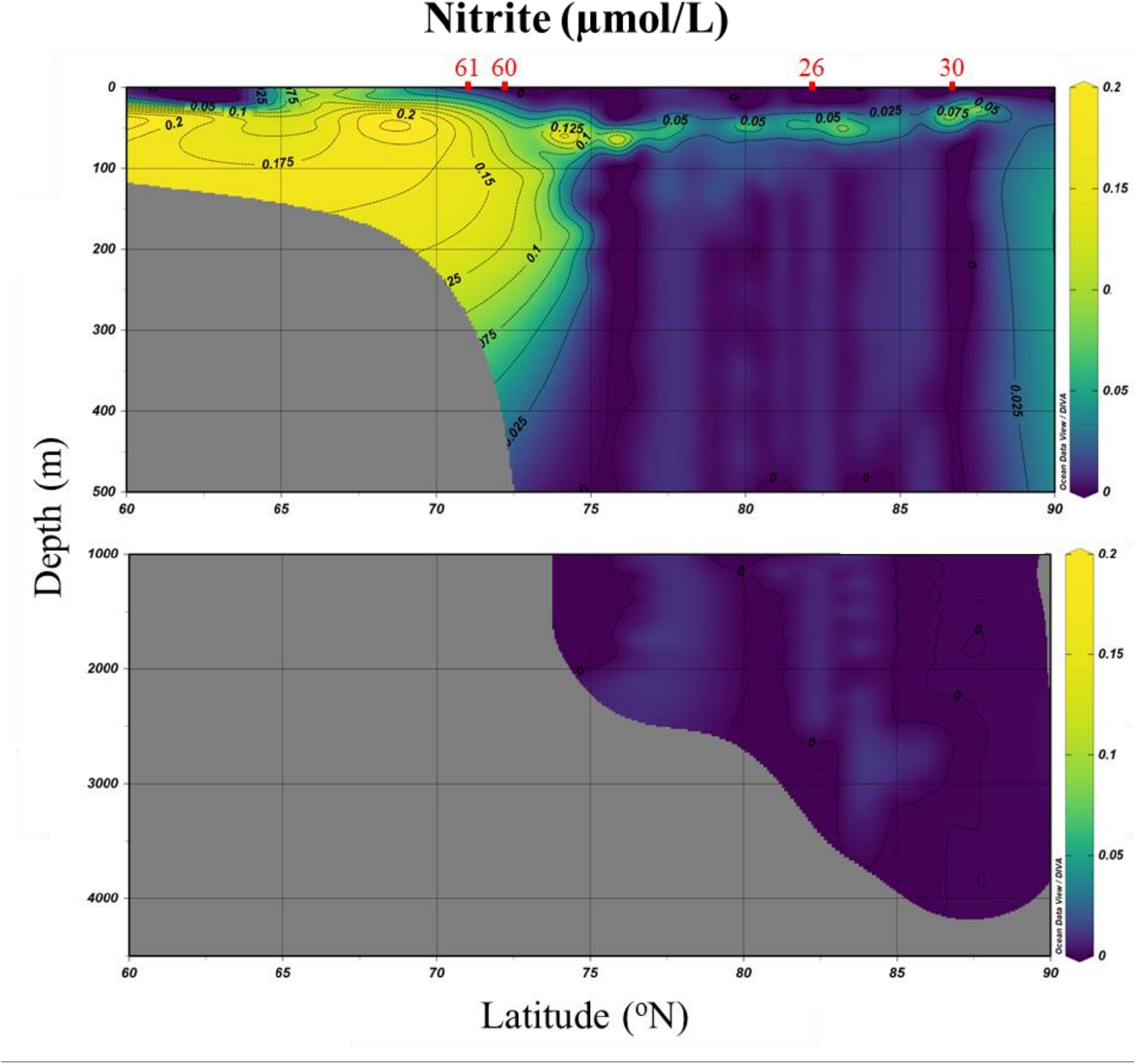
Section plot of nitrite concentrations for the Western Arctic Ocean during the GEOTRACES 2015 expedition (GN01). Values were gridded using weighted-average gridding in ODV. Metagenomic sampling data was collected at stations listed at the top (red).

### 3.2 Archaeal MAG identification

Metagenomic data from stations 26, 30, 60, and 61 were used for the identification of archaeal taxa as described in Sections 2.1–2.3. A single co-assembly of the 128 samples was produced with MetaSPAdes, resulting in 2,735,798 contigs. Contigs smaller than 1500 bp were removed, resulting in 224,116 contigs (867,388,435 bp) that were used for downstream processing. Three metagenomic binning algorithms (MetaBAT2; MetaBinner; MaxBin2) were used to identify bins that were consolidated using DAS Tool. A total of 155 bins with an average size of 2,312,350 bp (range: 211,901 to 14,059,051 bp) were recovered. Using GTDB-Tk for taxonomic assignment, 17 of the 155 bins were determined to be of archaeal origin (Table 2). Bins were identified in each of the three marine groups, although MG-II (Poseidoniales) contained the greatest amount of richness (11 bins). Bin quality was measured using CheckM (Table 2). CheckM completeness and contamination scores of > 90% and < 5% (“high-quality draft” as defined by the MIMAG standard; Bowers et al., 2017), were only met by bin 35_sub and bin 29, the remainder were classified as “medium-quality drafts” (> 50% completion, < 10% contamination), “low-quality drafts” (< 50% complete, < 10% contamination), or simply drafts (all others). The BV-BRC Comprehensive Genome Analysis tool was used to annotate the bins (Table 3). Bin 54 (MG-II) contained the fewest contigs (73), while bin 105_sub (MG-III) and bin 108 (MG-II) both contained the greatest number of contigs (530). The average genome size was 1,470,528 bp, which was within the normal size range for archaeal genomes. The smallest genomes were observed in the MG-I (maxbin.074_sub (908,512 bp); 166_sub (898,496 bp); 29 (956,250 bp)), while the largest was observed in MG-II (maxbin.078_sub (2,434,869 bp)). Contig N50 and L50 values were also calculated as a measure of bin quality. Bin 35_sub, bin 54, and bin 63 displayed the most desirable values (N50 > 20,000; L50 < 20) which were fairly consistent with the quality scores from CheckM. Lower quality bins determined by CheckM (145_sub, 161, and 166_sub) consequently displayed poorer N50 and L50 scores (N50 < 4000; L50 > 100 contigs). The number of coding sequences (CDSs) ranged from 1090 (bin 29) to 2395 (bin maxbin.078_sub), consistent with the genome sizes mentioned above. The lowest genomic GC percentages were observed in the MG-I MAGs (maxbin.074_sub (30.93%); 166_sub (33.31%); maxbin.015_sub (34.70%); 29 (34.73%)) but are well within the expected bounds of microbial GC content, which is typically between 30–50% GC. Both MG-III bins (35_sub (64.23%); 105_sub (60.59%)) had very high GC content, alongside members of MG-II (55_sub (59.05%); 54 (59.80%); maxbin.001_sub (55.96%); 63 (55.33%); 118 (59.81%); 108 (60.59%)). Phylogenetic analysis was performed on the archaeal bins (Figure 11). The phylogenetic analysis was consistent with the taxonomic assignments for all MAGs.

**Figure 11:**
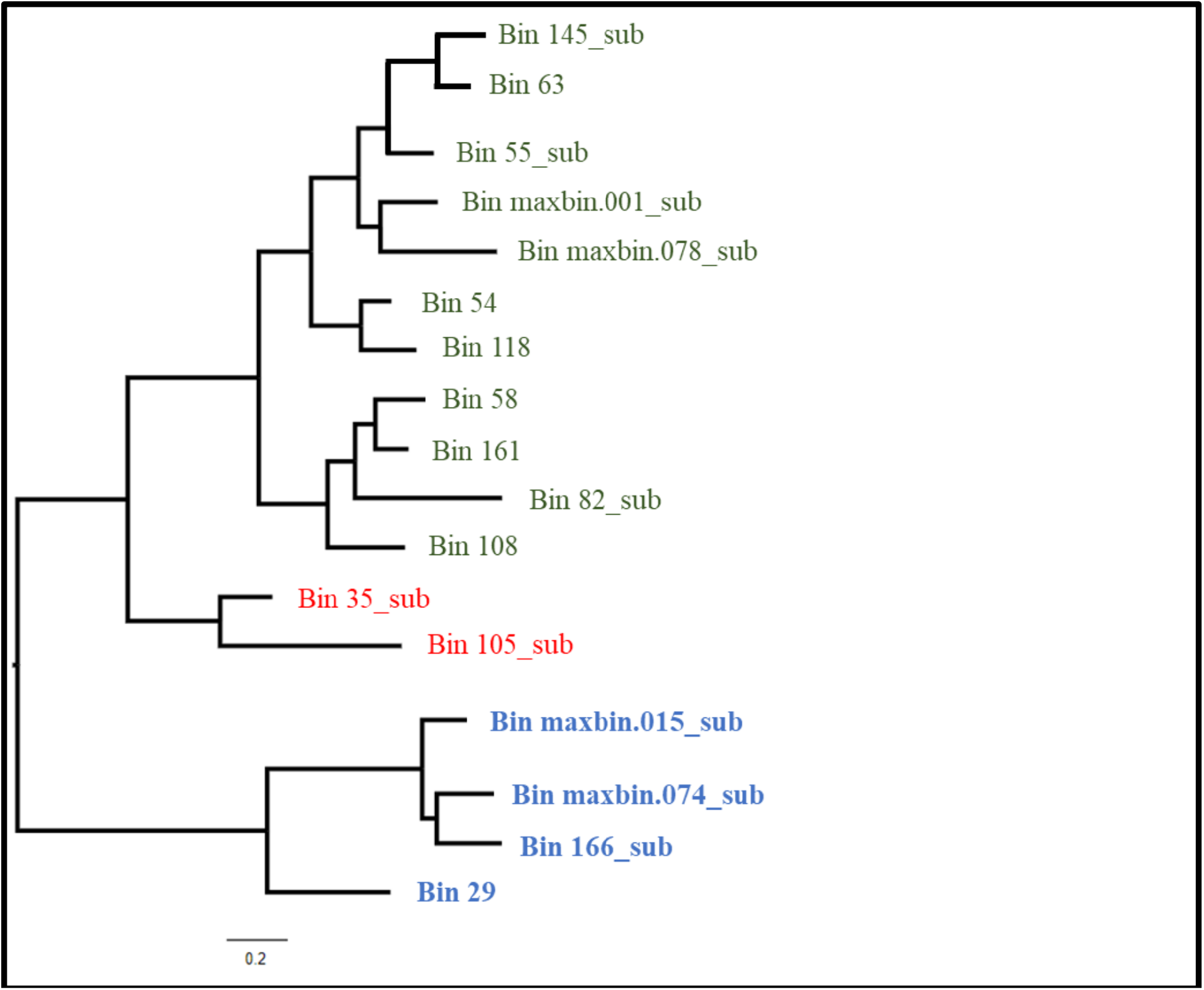
Phylogenetic tree of the archaeal MAGs identified in the Western Arctic Ocean during the 2015 GEOTRACES expedition (GN01). Clades are identified as MG-I (blue), MG-II (green), and MG-III (red).

**Table 2:** CheckM completeness and contamination estimates for the archaeal MAGs.

| Marine group | Bin ID | Completeness (%) | Contamination (%) |
|---|---|---|---|
| MG-III | 35_sub | 93.56 | 1.07 |
|  | 105_sub | 74.81 | 9.6 |
| MG-II | maxbin.078_sub | 84.36 | 4.4 |
|  | 55_sub | 82.49 | 0 |
|  | 54 | 82.36 | 0.8 |
|  | maxbin.001_sub | 82.23 | 2.46 |
|  | 63 | 81.24 | 0 |
|  | 58 | 79.33 | 0 |
|  | 82_sub | 78.95 | 1.93 |
|  | 118 | 73.92 | 5.65 |
|  | 108 | 63.88 | 1.92 |
|  | 145_sub | 59.05 | 3.6 |
|  | 161 | 50.92 | 1.85 |
| MG-I | 29 | 93.69 | 0.32 |
|  | maxbin.015_sub | 76.38 | 20.49 |
|  | maxbin.074_sub | 71.77 | 13.79 |
|  | 166_sub | 57.58 | 6.3 |

**Table 3:** Characteristics of the archaeal MAGs recovered from the 2015 GEOTRACES cruise, as reported by the BV-BRC whole-genome analysis tool. CDS, coding sequences.

| Bin | Genome length (bp) | GC (%) | Contigs | Contig N50 | Contig L50 | CDS |
|---|---|---|---|---|---|---|
| 35_sub | 1292743 | 64.23 | 117 | 28161 | 13 | 1309 |
| 105_sub | 1196390 | 60.59 | 530 | 2914 | 173 | 1623 |
| maxbin.078_sub | 2434869 | 44.39 | 405 | 18006 | 23 | 2395 |
| 55_sub | 1364045 | 59.05 | 110 | 18854 | 23 | 1256 |
| 54 | 1857474 | 59.80 | 73 | 36416 | 17 | 1561 |
| maxbin.001_sub | 1541469 | 55.96 | 152 | 21754 | 22 | 1465 |
| 63 | 1400658 | 55.33 | 133 | 22859 | 17 | 1277 |
| 58 | 1775200 | 52.89 | 113 | 22387 | 25 | 1535 |
| 82_sub | 1956988 | 44.54 | 217 | 13314 | 43 | 1757 |
| 118 | 1886358 | 59.81 | 501 | 4367 | 140 | 1961 |
| 108 | 1486156 | 60.59 | 530 | 2914 | 173 | 1623 |
| 145_sub | 1362659 | 56.59 | 422 | 3500 | 123 | 1412 |
| 161 | 1508422 | 58.12 | 522 | 3005 | 163 | 1734 |
| 29 | 956250 | 34.73 | 104 | 13853 | 21 | 1090 |
| maxbin.015_sub | 1172288 | 34.70 | 442 | 2698 | 147 | 1625 |
| maxbin.074_sub | 908512 | 30.93 | 326 | 2912 | 103 | 1360 |
| 166_sub | 898496 | 33.31 | 366 | 2566 | 125 | 1263 |

### 3.3 Archaeal relative abundance at Stations 26 and 30

To better understand abundance trends as a function of depth, abundance data for all bins was analyzed at stations 26 and 30. These stations were chosen for further analysis because archaeal relative abundance was high at these two stations, the water columns were deep, and the clearest distinction of water masses was observed here. The temperature minimums were observed at *∼*100 m for both stations (station 26: *−*1.37 °C; station 30: *−*1.44 °C) which marked the bottom of the PML (Figure 12). The temperature increased throughout the AW layer to a maximum at 450 m (*∼*1 °C), which began to decrease again moving into the BSB and ABW. Salinity was low at the surface (29 PSU) and increased until it reached a maximum at 300 m (*∼*35 PSU). Nitrite (NO_2_^−^) concentrations were much lower than nitrate (NO_3_^−^) concentrations throughout the depth profile (Figure 12). The relationship between the two was inversely proportional, where nitrite was higher at the surface and decreased with depth, and nitrate was lower at the surface and increased with depth. Nitrate concentrations reached a maximum at approximately 500 m for both stations (11.91 µmol L^−1^, Station 26; 12.37 µmol L^−1^, Station 30). Nitrite concentrations became undetectable below *∼*500 m for both stations. Clustering based on relative abundance patterns of all MAGs (155) uncovered three distinct communities based on depth (Figure 13). Using the water mass determinations from Section 3.1, the sample clusters from different depths could be associated with a water mass: PML, AW, and ABW. The ABW cluster consisted of samples below *∼*1000 m. The exceptions were samples ARODd.30.8 and RjKKY.30.8 both collected from a depth of 226 m. The AW cluster contained samples from depths of *∼*200 m to *∼*1100 m. The FSB and BSB can be seen within the AW cluster, separated by depths between *∼*200 to *∼*400 m and *∼*500 to *∼*1000 m, respectively. The final cluster, the PML, contained samples from *∼*25 m to *∼*200 m. It is important to note that these water mass distinctions are not rigid. Relative abundance was generally higher in the PML than the other layers for most bins, as was expected with the presence of phytoplankton at the surface. The bins were hierarchically clustered and divided into four clusters for analysis (Figure 13). Most bins (35_sub, 105_sub, maxbin.078_sub, 63, 54, 58, 82_sub, 29, maxbin.074_sub, and 166_sub) clustered into cluster A, with bins that displayed a higher relative abundance in the PML and AW layers. Bin 55_sub was found in cluster B which contained bins that displayed a mixed pattern of relative abundance across the samples. The final clusters (C and D) contained bins (maxbin.001_sub, 108, 118, 145_sub, 161, and maxbin.015_sub) with higher abundance in the ABW. Bin 118 was the only bin to cluster into cluster C which showed relatively high abundances at the very deep samples as well as some of the shallower samples collected. Bins from within the same phylogenetic lineage did not display similar abundance patterns at these two stations despite previously established distribution trends for the archaeal marine groups (Ionescu et al., 2009; Pereira et al., 2019; Rinke et al., 2019; Yilmaz et al., 2016). Overall, archaeal relative abundance at these stations appeared to increase with depth (Figure 14A). The MAGs were also analyzed by the proportion of total archaeal relative abundance (Figure 14B). The MG-I MAGs made up the largest proportion, up to 80% of the community, in the upper 400 m. The MG-II MAGs made up the largest proportion at most depths, particularly below 500 m. The MG-III MAG abundance proportion was fairly consistent throughout the water column (approximately 10% of the archaeal community). Relative abundances of each MAG with respect to the overall microbial community were also investigated. It was observed that the MG-I bins were found in greater abundances in the upper 500 m (Figure 15A). The MG-II varied greatly in their overall abundance but did increase with depth (Figure 15B). Bin 82_sub was the most abundant from this group. MG-III were much more abundant below 800 m dominated by bin 35_sub (Figure 15C), however their overall relative abundances were much lower than the other two marine groups.

**Figure 12:**
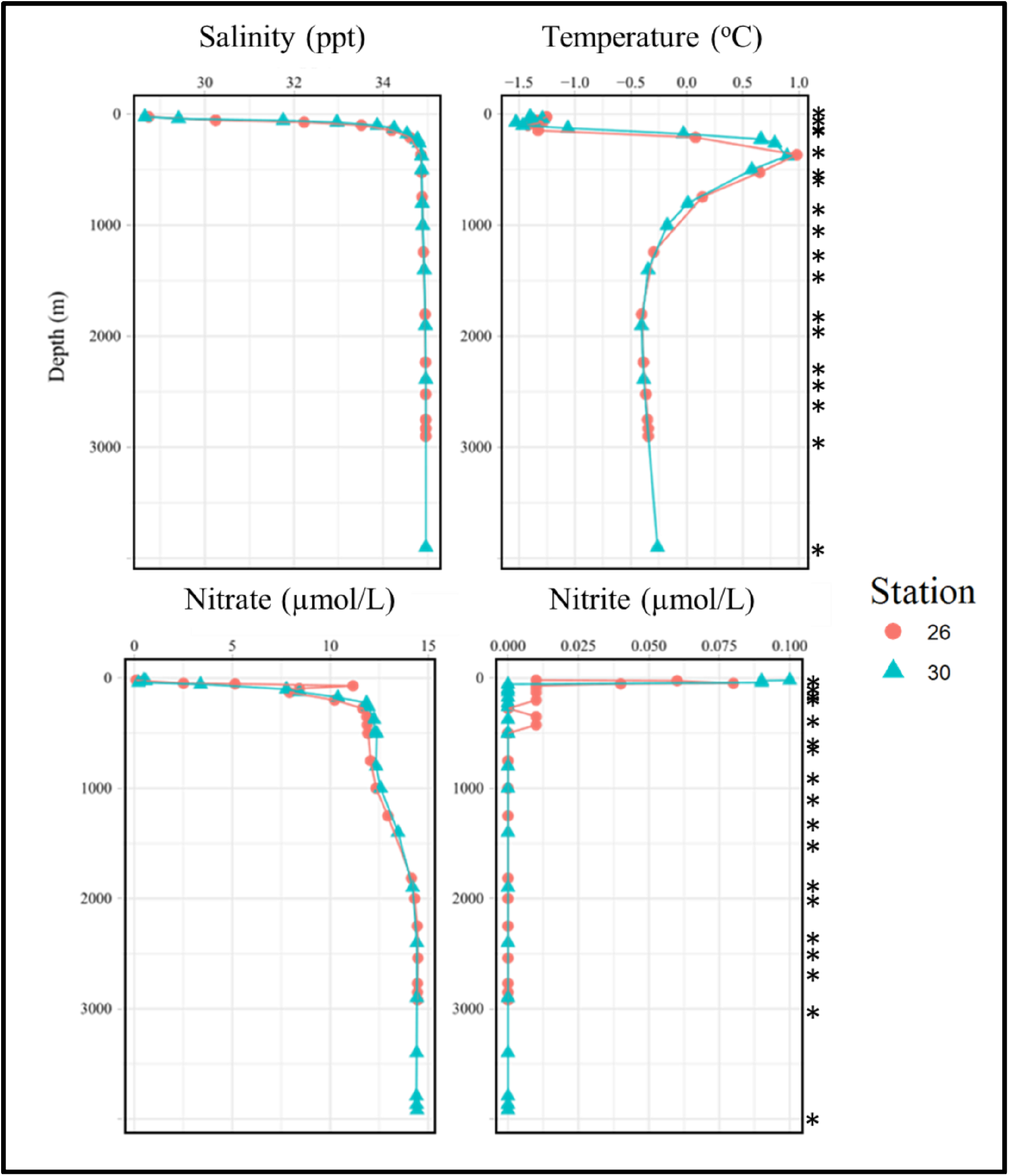
Depth profiles for salinity, temperature, nitrate, and nitrite from the 2015 GEOTRACES expedition from stations 26 (red circles) and 30 (blue triangles). The depths of metagenomic sampling are indicated on the right (*).

**Figure 13:**
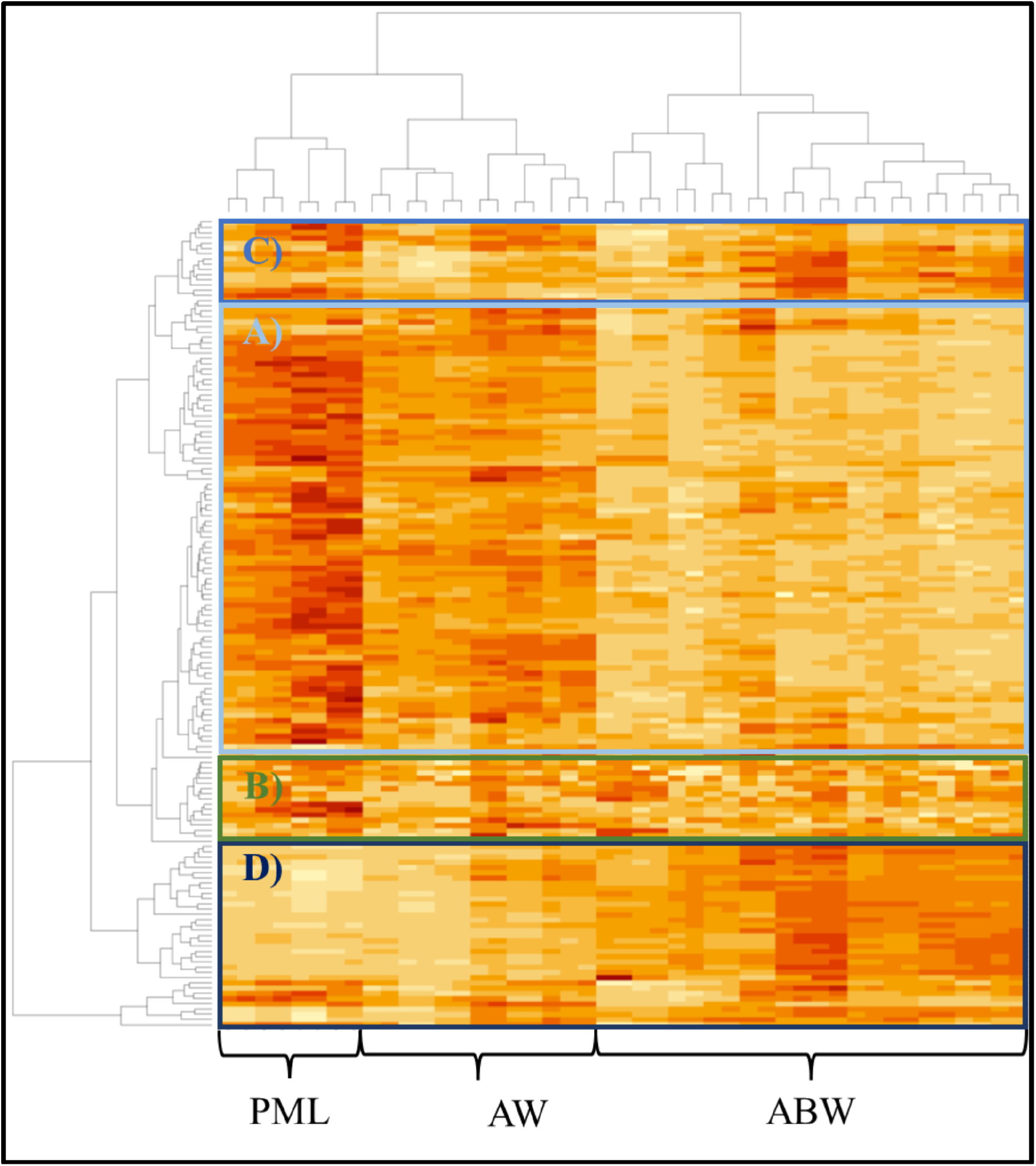
Heat map displaying abundance-based clustering of each bin (y-axis) and each sample (x-axis) at Stations 26 and 30 from the GEOTRACES 2015 expedition (GN01). Samples formed three main clusters based on depth: Arctic Ocean Basin water (ABW); Atlantic water (AW); Polar Mixed Layer (PML).

**Figure 14:**
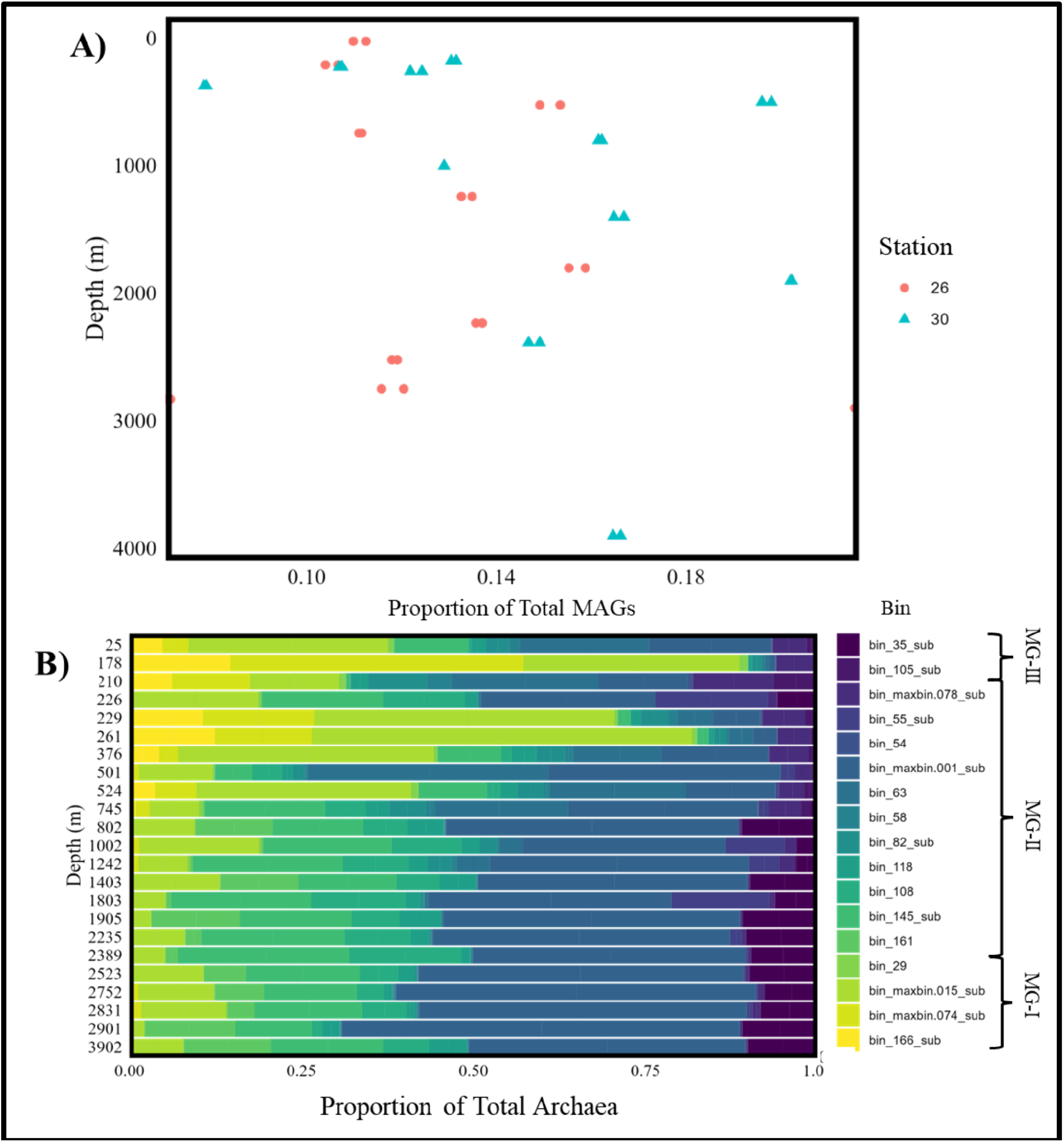
Relative abundance measurements for Archaea in the Western Arctic Ocean: A) total archaeal relative abundance as a proportion of total MAG abundance by depth; B) proportion of total archaeal abundance for all bins by depth.

**Figure 15:**
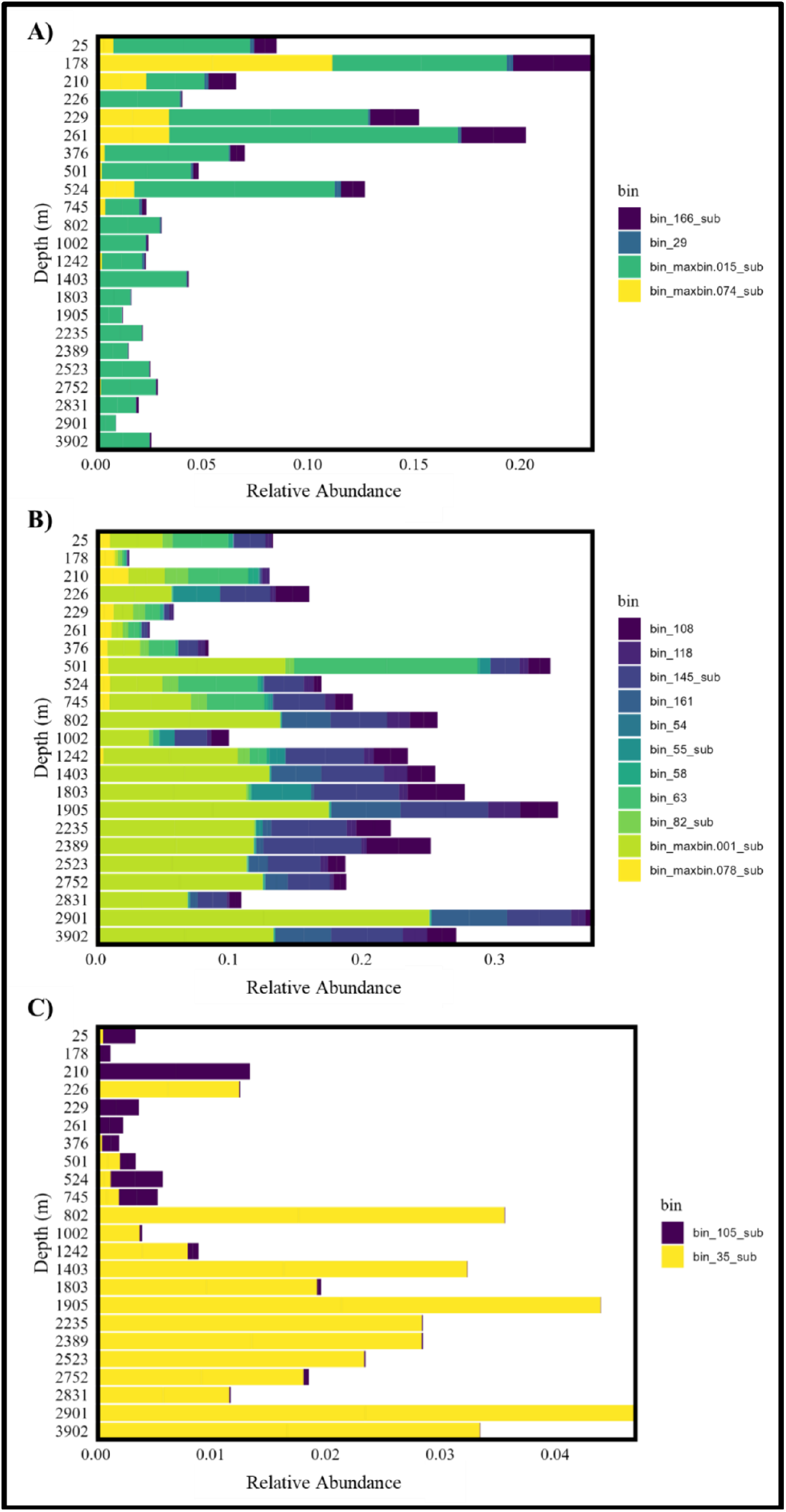
Archaeal relative abundance by depth for MG-I (A), MG-II (B), and MG-III (C) at station 26 and 30 from the 2015 GEOTRACES expedition (GN01).

### 3.4 Archaeal biogeography in the Western Arctic Ocean

To further explore the archaeal distribution in the Western Arctic Ocean, the relative abundance of each bin was plotted by latitude and depth across a dataset of several hundred samples. Three main abundance distribution patterns were observed, which were variable within each marine group (Figures 16–20). These three abundance distribution ecotypes corresponded with the water masses determined in Section 3.1: PML, AW, and ABW. Again, these descriptions of water mass layers are not definitive, and thus the ecotypes were simplified to surface, mid-depth, and deep ecotypes. The MG-I Archaea demonstrated surface and deep distributions (Figure 16). In fact, maxbin.074_sub had the highest relative abundance of the four bins and was found in greater amounts in the surface layers just north of the coast (Figure 16A). The abundances observed did correspond with regions with elevated nitrite concentrations (Figure 10). It was observed that the MG-II Archaea displayed all three distribution ecotypes: Bins 54 and maxbin.078_sub displayed a surface ecotype (Figure 17); Bins 58, 118, 82_sub, and 63 displayed a mid-depth ecotype (Figure 18); and Bins 55_sub, maxbin.001_sub, 161, 108, and 145_sub displayed a deep ecotype (Figure 19). Bin maxbin.078_sub was very abundant at the surface and was found at lower latitudes where water temperatures were much higher, as was expected. The mid-depth MG-II were lower in their abundances overall compared to maxbin.078_sub. In contrast, maxbin.001_sub exhibited comparable abundance measurements to the surface residing maxbin.078_sub while presenting a deep ecotype. The MG-III Archaea displayed both surface (bin 105_sub; Figure 20A) and deep (35_sub; Figure 20B) ecotypes. Interestingly, bin 105_sub, although surface residing, did display a different pattern of distribution where the abundance was greatest closer to the pole rather than the shelf. The relative abundance distributions were compared to the phylogenetic tree structure (Figure 11) to see if the surface, mid-depth, and deep ecotypes were more closely related to one another. There was no consistency between the phylogenetic structure and the water profile ecotypes displayed by the MAGs. For example, maxbin.074_sub which was identified as a surface residing MG-I was not less related to the three other deep residing MG-I MAGs. The majority of the associated ecotypes were consistent with the relative abundance clustering that was seen at stations 26 and 30 (Figure 13). There were three exceptions. Bin maxbin.015_sub (MG-I) clustered in Group D at stations 26 and 30 which were associated with higher relative abundances from the deeper water samples. Although this bin was determined to have a mid-depth ecotype, the abundances were quite high at 800–1000 m which roughly marks the bottom of the AW layer. This group was also found at moderate abundances in the ABW. The other two bins that had discrepancies were MG-II MAGs. Bin 118 was determined to have a mid-depth ecotype, however it clustered with Group C, which was associated with higher abundances at greater depths as well as higher abundances in some of the shallower samples. Bin 55_sub was identified as belonging to Group B, which displayed a mixed abundance pattern at the two stations. Although it was characterized as having a deep ecotype, this mixed abundance behaviour was also seen in the biogeography as abundance was high in the 200 m range and 2000 m range. The addition of more southern stations (60 and 61) has elucidated the biogeography of these organisms that would have been missed with fewer stations.

**Figure 16:**
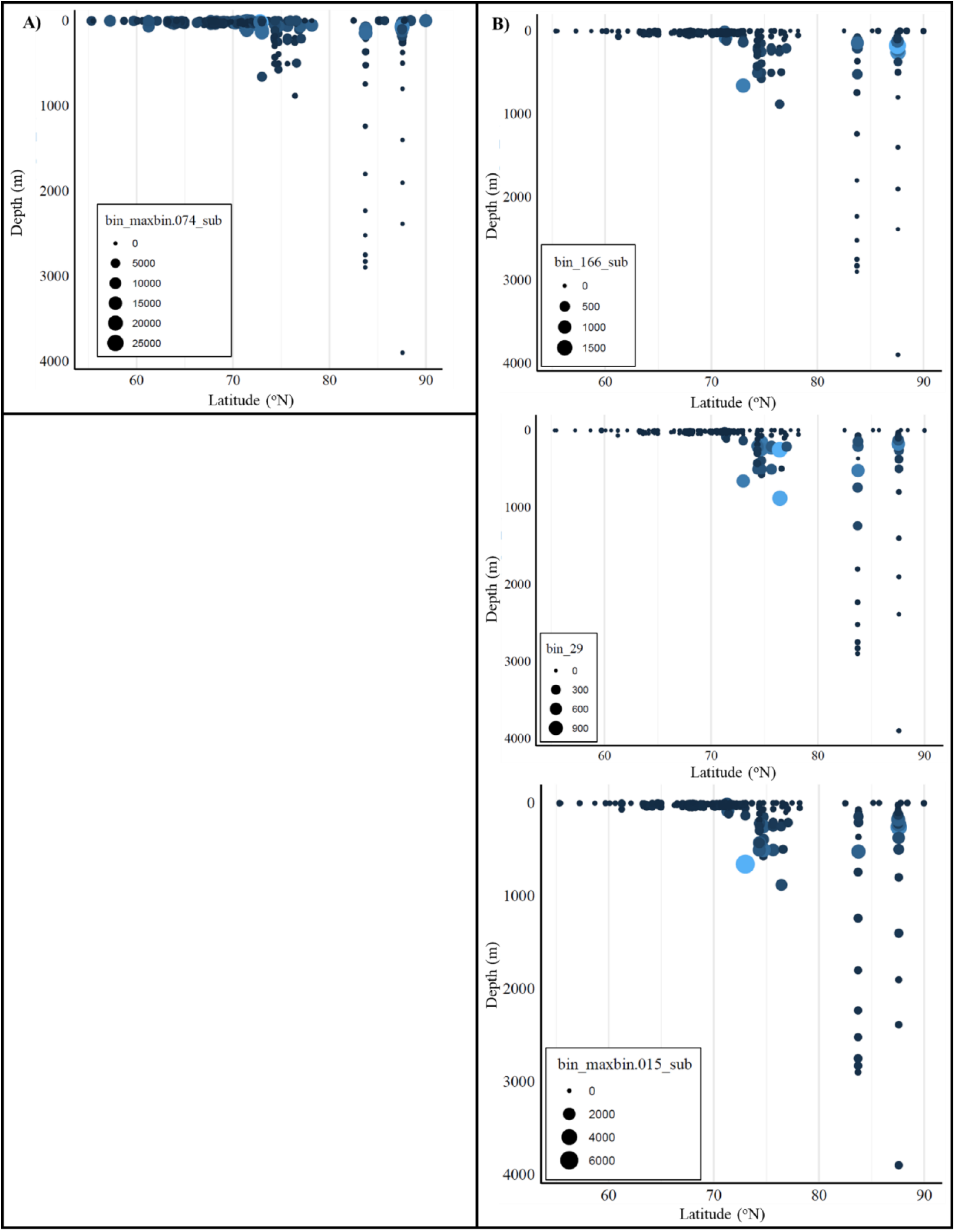
Relative abundance distributions (*×*10^6^) of the MG-I archaeal MAGs identified in the Western Arctic Ocean displaying a surface ecotype (A) and a mid-depth ecotype (B). Colours correspond with the abundances (light blue is higher relative abundance and dark blue is lower relative abundance).

**Figure 17:**
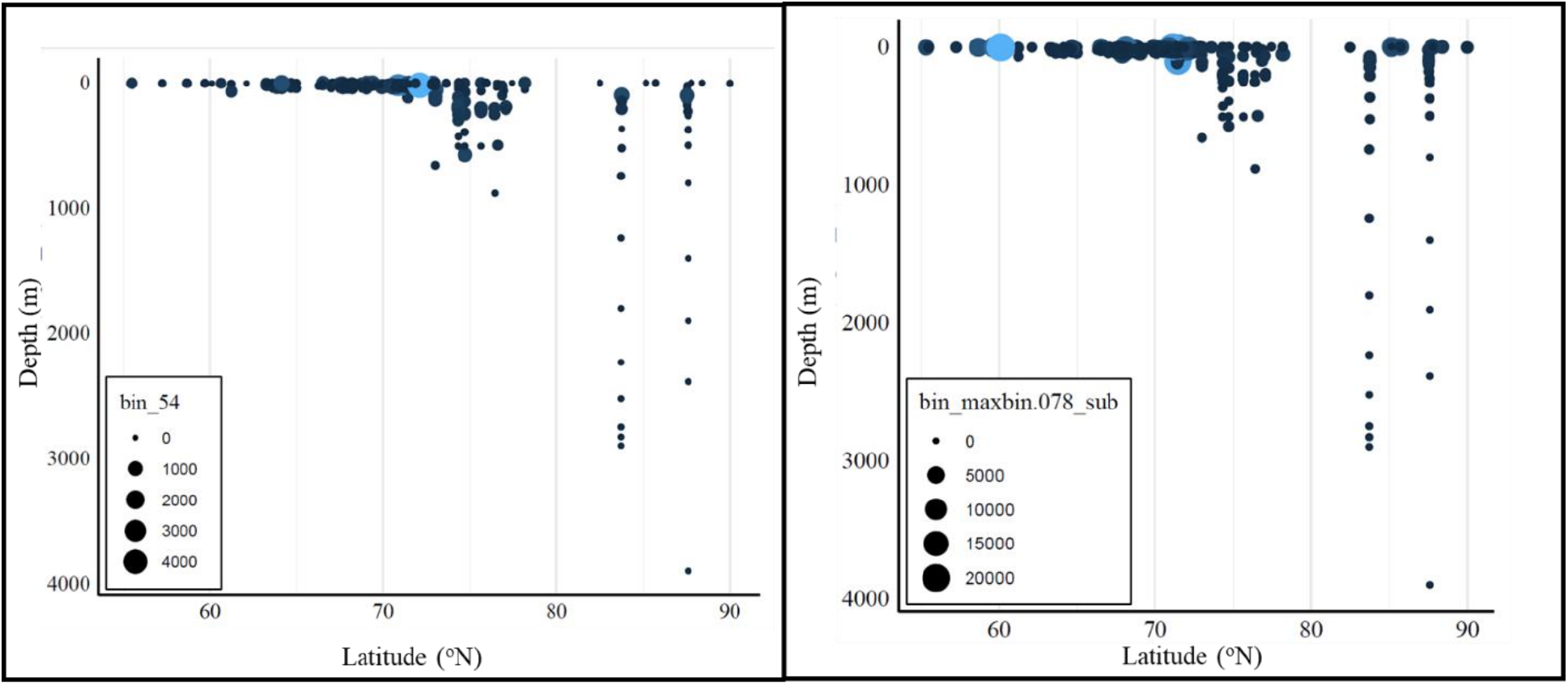
Relative abundance distributions (*×*10^6^) of the MG-II archaeal MAGs identified in the Western Arctic Ocean displaying a surface ecotype. Colours correspond with the abundances (light blue is higher relative abundance and dark blue is lower relative abundance).

**Figure 18:**
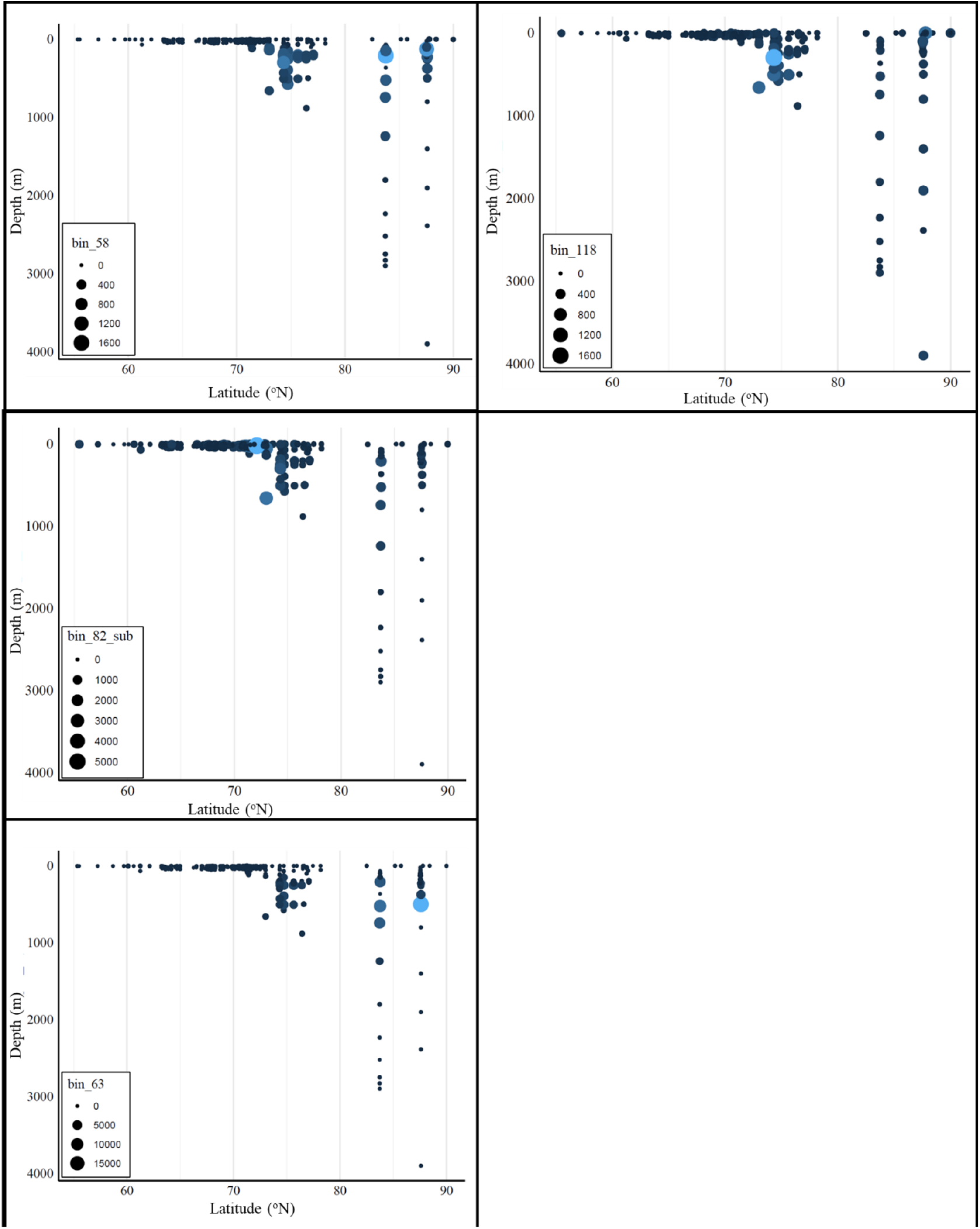
Relative abundance distributions (*×*10^6^) of the MG-II archaeal MAGs identified in the Western Arctic Ocean displaying a mid-depth ecotype. Colours correspond with the abundances (light blue is higher relative abundance and dark blue is lower relative abundance).

**Figure 19:**
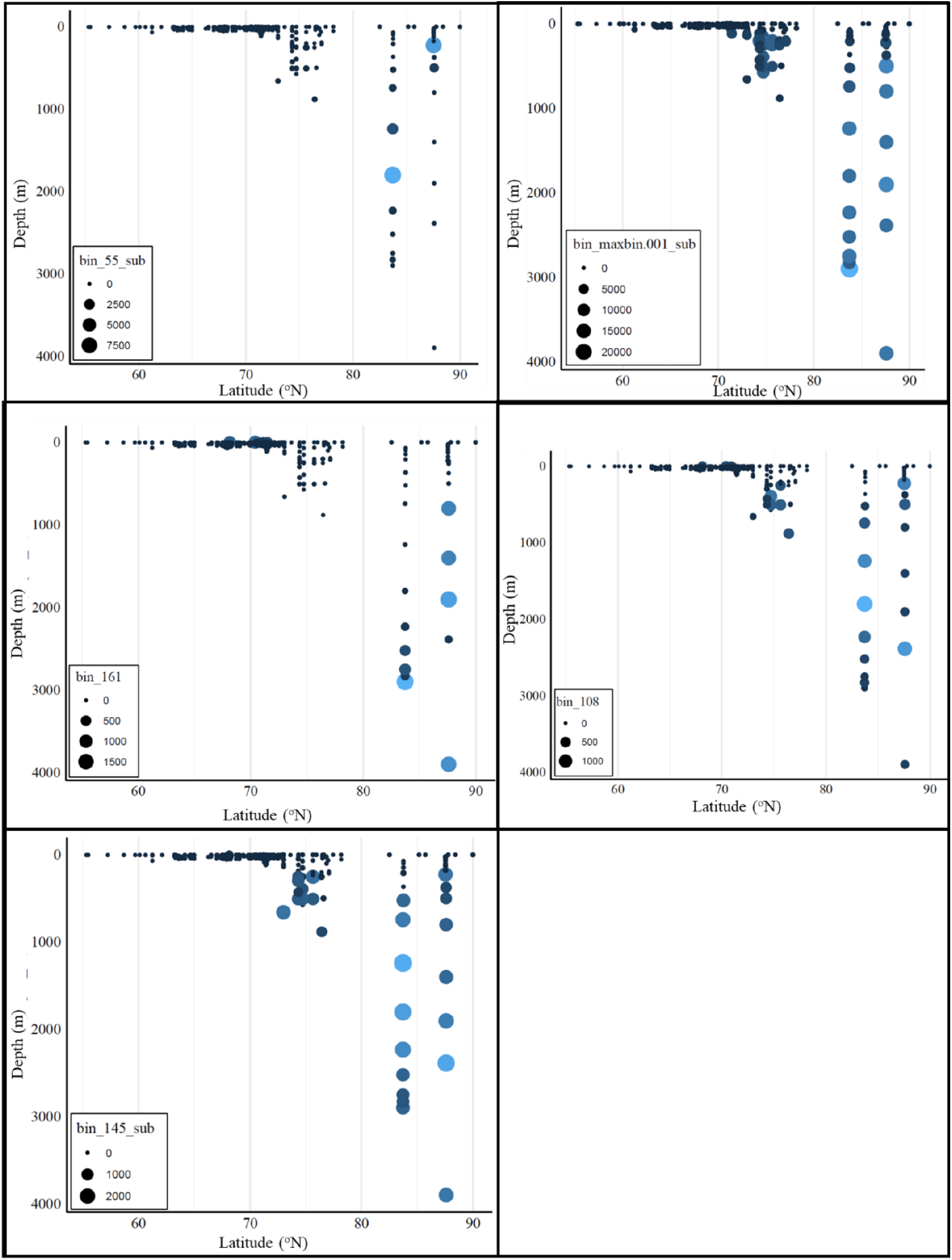
Relative abundance distributions (*×*10^6^) of the MG-II archaeal MAGs identified in the Western Arctic Ocean displaying a deep ecotype. Colours correspond with the abundances (light blue is higher relative abundance and dark blue is lower relative abundance).

**Figure 20:**
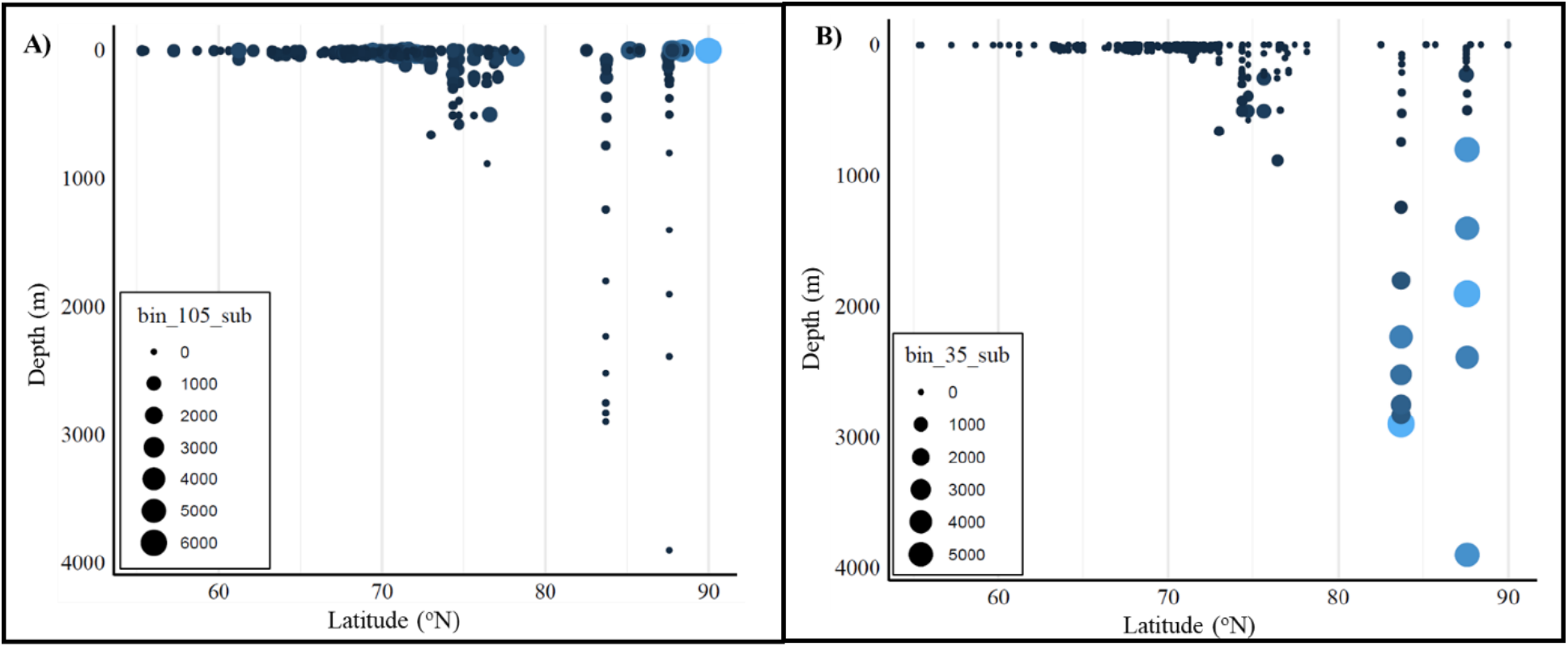
Relative abundance distributions (*×*10^6^) of the MG-III archaeal MAGs identified in the Western Arctic Ocean displaying a surface ecotype (A) and a deep ecotype (B).

## 4 Discussion

### 4.1 Archaeal MAG identification and the use of metagenomic analysis

Metagenomics is a relatively new technique for the study of environmental microbial samples. Historically, organisms were classified using 16S rRNA sequencing, also referred to as tag sequencing. The general process of tag sequencing involves sequencing a small region of the genome, typically the 16S rRNA region, selected with the use of primers for taxonomic assignment. There are three major drawbacks to this type of analysis. The first is that the primers chosen for the sequencing process can greatly affect the downstream results. There are many aspects to consider when choosing primer sets: the primer length, the specificity, and other physical and chemical properties that can affect the efficacy of the analysis. The chosen primers therefore have potential to completely miss entire clades of organisms, especially those that are rare and relatively unique. The second drawback is that the 16S rRNA region is fairly short, relative to the entire genome, and may not be effective in elucidating very closely related organisms, resulting in incorrect taxonomic assignments. The final drawback of this technique is that no functional information is provided. Therefore, the biogeochemical processes being performed in that respective environment are left unknown. Metagenomic analysis attempts to resolve these issues by removing the need to use primers and by sequencing all regions of the genome to obtain a higher level of phylogenetic resolution and provide functional information. In this study, 17 archaeal MAGs were identified in the Western Arctic Ocean. Although relative abundances and diversity were low, members of all three marine groups were identified. They displayed ranges in completeness and contamination, although most were categorized as medium- or high-quality drafts. Both MG-III bins (35_sub (64.23%); 105_sub (60.59%)) had very high GC content, alongside members of MG-II (55_sub (59.05%); 54 (59.80%); maxbin.001_sub (55.96%); 63 (55.33%); 118 (59.81%); 108 (60.59%)) (Table 3). GC content, the proportion of guanine and cytosine in the DNA, is often used to infer the level of DNA stability within a genome as guanines and cytosines can form three hydrogen bonds. High GC content is often associated with microorganisms living in harsh environments such as those with high temperatures, high salt concentrations, or high levels of predation. These observations are not typically associated with organisms observed in colder habitats. There were discrepancies observed between previous studies of Archaea in marine environments and this study when exploring the spatial distributions of the archaeal marine groups. It is likely that these differences were a result of using different sequencing technologies. The most probable cause for these differences is that 16S rRNA sequencing causes closely related organisms to be identified as the same lineage, meaning that the higher abundance organisms would display a stronger genomic signal, therefore biasing the results of where these organisms were found. The use of metagenomic sequencing has allowed for the separation of these closely related genomes into individual bins so that the diversity of distribution patterns could be observed.

### 4.2 Archaeal distributions are influenced by water masses rather than strictly by light penetration and temperature

The Arctic Ocean is a small, enclosed basin that provides a harsh environment for microbial life. The temperatures are low (< 20 °C), the salinities are high (> 20 ppt), and light is variable with season and depth. This ocean offers a unique look at the microbial communities residing there, their biogeography, as well as their biogeochemical processes. The water masses of the Western Arctic Ocean were defined by their thermocline and nutrient properties. These water mass layers were simplified to contain the Summer Surface Layer, Polar Mixed Layer, Atlantic Water layer, and Arctic Ocean Basin Water layer. The overall microbial communities were shown to cluster into three major groups (Figure 13). These groups were differentiated based on the water mass layer of collection. It was shown here that the different water mass layers harbour distinct and unique communities. Previous studies (Ionescu et al., 2009; Liu et al., 2018; Pereira et al., 2019; Rinke et al., 2019) have observed that the three archaeal marine groups have specific and consistent distributions within a marine water column. For example, the MG-I are commonly found to be the most abundant archaeal clade in marine environments overall but are generally found at higher abundances at or below 200 m (Baer et al., 2014; Christman et al., 2011). The MG-II are often found at higher abundances in the upper 200 m, however are present at lower abundances at greater depths (Liu et al., 2017). Finally, the MG-III are often found at very low abundances fairly deep within the water column. It has been suggested that these variations in distributions are controlled by microbial responses to light penetration into the water, and secondly by temperature. When analyzing the individual bins from each marine group from all sampling locations and depths it was shown that these results differed from previous studies. Here it was observed that MAGs from the different archaeal marine groups did not show a collective preference for any one water mass (Figures 16–20). Bins from within the same phylogenetic lineage did not display similar abundance patterns despite previously established distribution trends for the archaeal marine groups (Ionescu et al., 2009; Pereira et al., 2019; Rinke et al., 2019; Yilmaz et al., 2016). There was a considerable amount of variation within the groups themselves, suggesting that there was niche overlap due to shared environmental preferences and metabolic interactions involving the exchange of nutrients. Many of the previous studies that have looked at archaeal biogeography in the oceans have claimed that the phylogenetic organization of the groups influences the types of niches they reside in and that these characteristics are consistent and specific to each clade of Archaea. However, it was seen here that the type of biogeographical distributions observed were not specific to any one clade. All three marine groups identified in the Western Arctic had multiple biogeographical distributions. Although the temperature and light penetration are connected to the different water mass layers, these parameters do not affect the distributions of the entire clade in the same way. The distributions of the three marine groups were investigated at Stations 26 and 30. Both of these stations were located above 80 °N, closer to the pole. Nitrate concentrations at these stations followed a classic nutrient profile where concentrations were low at the surface and increased with depth (Figure 12). Low levels of nitrate are indicative of nutrient utilization by phyto- and picoplankton, consistent with the expected surface blooms. At these stations, it was observed that nitrite concentrations were elevated at the surface (Figure 12) which is often associated with a higher prevalence of MG-I activity (Liu et al., 2018) because of their ability to oxidize ammonia to nitrite. We did see a moderately elevated MG-I relative abundance at 25 m (Figure 15A); however, it is difficult to correlate these two findings because there are other ammonia-oxidizing microbes that can be influencing this observation, especially in late summer months when phytoplankton blooms are expected to be present. Other studies have observed MG-I at higher abundances below the thermocline (Ionescu et al., 2009), which has been used to suggest that MG-I members prefer an environment that is cooler. It was observed that the MG-I bins were found in greater abundances in the upper 500 m (Figure 15A). There were two distribution ecotypes observed for the MG-I (Figure 16). This is broadly consistent with other studies that found MG-I abundances increase below 200 m (Church et al., 2003; Herfort et al., 2007; Liu et al., 2018; Pitcher et al., 2011; Reji et al., 2019; Wuchter et al., 2006). MG-II Archaea are generally thought to be surface residing, with many studies observing them at high abundances in the top 200 m of the water column (Pereira et al., 2019). The MG-IIb are commonly seen at depths around 200 m, but most studies did not sample below 500 m (Church et al., 2003; Offre et al., 2013; Pereira et al., 2019; Rinke et al., 2019). MG-II are usually the most abundant Archaea in surface waters, so it is unusual to see the MG-II MAGs dominating the community below the epipelagic zone (Figure 15B). This observation has two explanations. One, that the abundances truly are higher at depths greater than 500 m. As mentioned previously, many studies did not sample below 500 m, and therefore did not observe this finding. The second explanation involves the use of relative abundance as a measurement. The calculation of relative abundance involves mapping the reads from the sequencing to the assembled contigs. This provides a proportion that represents how many times that contig was observed in the sample. This can then be used to calculate how many times each bin was observed in the sample. Relative abundance is therefore a measure of the total proportion of microbial bins identified. Consequently, when an increase in relative abundance is detected, it could actually be that there were fewer organisms observed, rather than an increase in absolute counts. The MG-II displayed all three distribution ecotypes (Figures 17–19). The group of deep residing MG-II could be a new subgroup that has not yet been defined. This discrepancy with previous studies is likely due to a lack of sampling depth in previous studies. Although previously thought of as surface residing organisms, it is clear now that as a phylogenetic group they are not all surface residing and that their biogeography is much broader than previously stated. MG-III are among the least understood members of marine microbial communities. They are typically considered low-abundance members of deep-sea environments (Galand et al., 2010). The MG-III MAG relative abundance was fairly consistent throughout the water column (approximately 10% of the archaeal community) (Figure 14B), which is consistent with the literature (Haro-Moreno et al., 2017) as these organisms are often identified at low abundances. Their relative abundances did increase with depth (Figure 15C), as was expected. It is important to note again that this observation could be due to the use of relative abundance measurement as well. The two MG-III MAGs displayed a surface and deep ecotype. A recent study found members of MG-III in both the photic zones and deep-sea samples (Haro-Moreno et al., 2017). Haro-Moreno et al. (2017) suggest that the epipelagic MG-III evolved from a deep-water (bathypelagic) ancestor as they contained many light-protecting proteins and pathways derived from bacterial origin, likely obtained through horizontal gene transfer. Beyond the environmental structuring observed here, there may also be biological structuring occurring where the taxa present can be predicted based on other taxa there. There are factors influencing communities to work together rather than as individuals. It can be helpful to think of the microbial community structure like a body of microorganisms that are working within a defined geographical space. The geographical space that holds these microbial bodies is connected to the water mass layers. The water masses are the current assumption influencing the distributions but there could be a deeper link within the community. Investigating the co-occurrence of archaeal taxa with other members of the community is likely to elucidate the biological factors that are influencing this community structure.

## 5 Conclusions

In this study it was hypothesized that the water masses would be the strongest factor influencing the archaeal relative abundance patterns in the Western Arctic Ocean. It was shown that the overall communities within the water mass layers were distinct. It was also shown that the archaeal marine groups did not show a consistent distribution pattern, like what had been previously observed. Through the analysis of the relative abundance distributions of the individual archaeal MAGs, this study expands our current understanding of archaeal marine biogeography. While it has been suggested that the three marine groups have specific and consistent distributions, we show here, with increased phylogenetic resolution, expanded sampling depths, expanded water mass characterizations, and the use of metagenomic tools, that bins within each marine group displayed variability in their abundance mapping, suggesting that the previous claim that these traits are specific and consistent does not hold in this system. To further understand the ecological context of Arctic Archaea, determining absolute abundance measurements is the first step. Additionally, further exploration of the environmental parameters associated with the different water masses that may be influencing the abundance distributions will be helpful. As was discussed above, there are also biological factors that influence microbial communities. Investigating the co-occurrence of different taxa in the environment would be the first step in understanding the biological interconnectedness.

## Acknowledgements

This manuscript is adapted from Chapter 2 of the MSc thesis of K. Green (University of Manitoba, 2023). We thank Drs. C. J. Mundy and Jose Luis Rodriguez-Gil for their guidance throughout this work. We thank Katlin Bowman, Kyle Dilliplaine, Pete Morton, and everyone aboard the USCGC *Healy* during the 2015 GEOTRACES cruise (HLY1502) who was involved in the collection of the metagenomic samples.

## Data availability

Raw metagenomic sequence data analyzed in this study are available from the International Nucleotide Sequence Database Collaboration (NCBI/ENA) under BioProject accessions PRJEB14154 (Arctic Ocean metagenomes from the 2015 USCGC *Healy* GEOTRACES cruise, HLY1502) and PRJNA681031 (Arctic metagenomes sequenced on the Illumina MiSeq).

## Author contributions

**Kari Green:** Formal analysis, Investigation, Data curation, Visualization, Writing – original draft. **R. Eric Collins:** Conceptualization, Methodology, Resources, Data curation, Supervision, Project administration, Funding acquisition, Writing – review & editing.

## Supplementary Information

**Figure S1:**
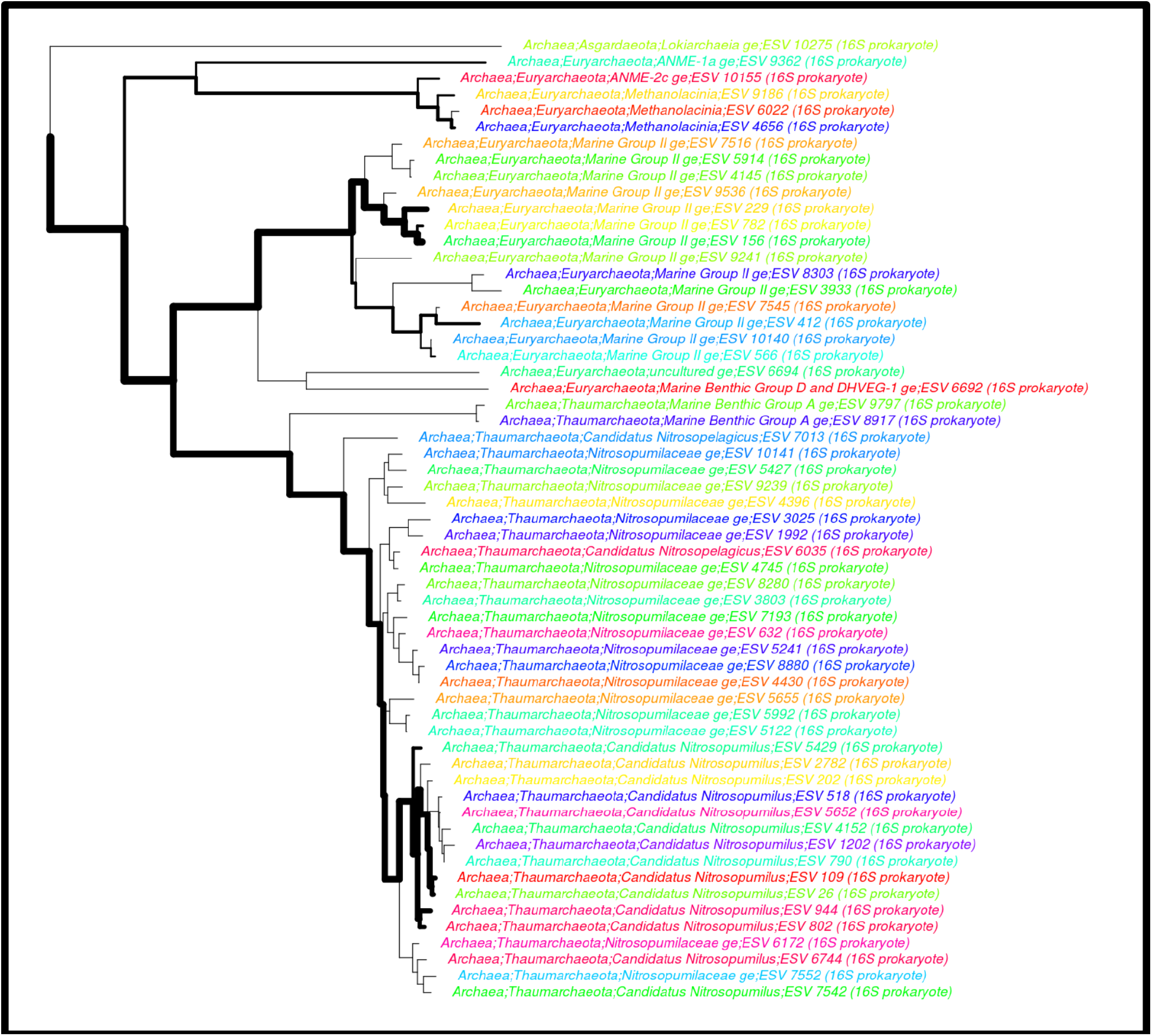
Phylogenetic tree of 16S rRNA gene sequencing data from samples collected in the Chukchi Sea by the Collins lab, plotted using Microscape (http://cryomics.org/microscape). Metagenomic data from a subset of this sample set were analyzed in this study. Each leaf on the tree is an exact sequence variant (ESV), corresponding approximately to the species or ecotype level. The line weights are proportional to the mean relative abundance of each ESV in the dataset. Branch lengths represent the relative evolutionary time between each node.

## References

Panagiotis S Adam, Guillaume Borrel, Céline Brochier-Armanet, and Simonetta Gribaldo. The growing tree of Archaea: new perspectives on their diversity, evolution and ecology. The ISME Journal, 11(11):2407–2425, August 2017. ISSN 1751-7370. doi: 10.1038/ismej.2017.122.

Nestor Arandia-Gorostidi, Anders K. Krabberød, Ramiro Logares, Ina Maria Deutschmann, Renate Scharek, Xosé Anxelu G. Morán, Felipe González, and Laura Alonso-Sáez. Novel Interactions Between Phytoplankton and Bacteria Shape Microbial Seasonal Dynamics in Coastal Ocean Waters. Frontiers in Marine Science, 9, July 2022. ISSN 2296-7745. doi: 10.3389/fmars.2022.901201.

Steven E. Baer, Tara L. Connelly, Rachel E. Sipler, Patricia L. Yager, and Deborah A. Bronk. Effect of temperature on rates of ammonium uptake and nitrification in the western coastal Arctic during winter, spring, and summer. Global Biogeochemical Cycles, 28(12):1455–1466, December 2014. ISSN 1944-9224. doi: 10.1002/2013gb004765.

Brett J. Baker, Valerie De Anda, Kiley W. Seitz, Nina Dombrowski, Alyson E. Santoro, and Karen G. Lloyd. Diversity, ecology and evolution of Archaea. Nature Microbiology, 5(7):887–900, May 2020. ISSN 2058-5276. doi: 10.1038/s41564-020-0715-z.

Derek M. Bickhart, Mikhail Kolmogorov, Elizabeth Tseng, Daniel M. Portik, Anton Korobeynikov, Ivan Tolstoganov, Gherman Uritskiy, Ivan Liachko, Shawn T. Sullivan, Sung Bong Shin, Alvah Zorea, Victòria Pascal Andreu, Kevin Panke-Buisse, Marnix H. Medema, Itzhak Mizrahi, Pavel A. Pevzner, and Timothy P. L. Smith. Generating lineage-resolved, complete metagenome-assembled genomes from complex microbial communities. Nature Biotechnology, 40(5):711–719, January 2022. ISSN 1546-1696. doi: 10.1038/s41587-021-01130-z.

Robert M. Bowers, Nikos C. Kyrpides, Ramunas Stepanauskas, et al. Minimum information about a single amplified genome (MISAG) and a metagenome-assembled genome (MIMAG) of bacteria and archaea. Nature Biotechnology, 35(8):725–731, 2017. doi: 10.1038/nbt.3893.

Jason E Box, William T Colgan, Torben Røjle Christensen, Niels Martin Schmidt, Magnus Lund, Frans-Jan W Parmentier, Ross Brown, Uma S Bhatt, Eugénie S Euskirchen, Vladimir E Romanovsky, John E Walsh, James E Overland, Muyin Wang, Robert W Corell, Walter N Meier, Bert Wouters, Sebastian Mernild, Johanna Mård, Janet Pawlak, and Morten Skovgård Olsen. Key indicators of Arctic climate change: 1971–2017. Environmental Research Letters, 14(4):045010, April 2019. ISSN 1748-9326. doi: 10.1088/1748-9326/aafc1b.

Thomas Brettin, James J. Davis, Terry Disz, Robert A. Edwards, Svetlana Gerdes, Gary J. Olsen, Robert Olson, Ross Overbeek, Bruce Parrello, Gordon D. Pusch, Maulik Shukla, James A. Thomason, Rick Stevens, Veronika Vonstein, Alice R. Wattam, and Fangfang Xia. RASTtk: A modular and extensible implementation of the RAST algorithm for building custom annotation pipelines and annotating batches of genomes. Scientific Reports, 5(1), February 2015. ISSN 2045-2322. doi: 10.1038/srep08365.

Céline Brochier-Armanet, Bastien Boussau, Simonetta Gribaldo, and Patrick Forterre. Mesophilic crenarchaeota: proposal for a third archaeal phylum, the Thaumarchaeota. Nature Reviews Microbiology, 6(3):245–252, March 2008. ISSN 1740-1534. doi: 10.1038/nrmicro1852.

Brian Bushnell, Jonathan Rood, and Esther Singer. BBMerge – Accurate paired shotgun read merging via overlap. PLOS ONE, 12(10):e0185056, October 2017. ISSN 1932-6203. doi: 10.1371/journal.pone.0185056.

Bonnie X. Chang and Allan H. Devol. Seasonal and spatial patterns of sedimentary denitrification rates in the Chukchi sea. Deep Sea Research Part II: Topical Studies in Oceanography, 56(17): 1339–1350, August 2009. ISSN 0967-0645. doi: 10.1016/j.dsr2.2008.10.024.

Pierre-Alain Chaumeil, Aaron J Mussig, Philip Hugenholtz, and Donovan H Parks. GTDB-Tk v2: memory friendly classification with the genome taxonomy database. Bioinformatics, 38(23): 5315–5316, October 2022. ISSN 1367-4811. doi: 10.1093/bioinformatics/btac672.

Lijing Cheng, Kevin E. Trenberth, John Fasullo, Tim Boyer, John Abraham, and Jiang Zhu. Improved estimates of ocean heat content from 1960 to 2015. Science Advances, 3(3), March 2017. ISSN 2375-2548. doi: 10.1126/sciadv.1601545.

Glenn D. Christman, Matthew T. Cottrell, Brian N. Popp, Elizabeth Gier, and David L. Kirchman. Abundance, Diversity, and Activity of Ammonia-Oxidizing Prokaryotes in the Coastal Arctic Ocean in Summer and Winter. Applied and Environmental Microbiology, 77(6):2026–2034, March 2011. ISSN 1098-5336. doi: 10.1128/aem.01907-10.

Matthew J. Church, Edward F. DeLong, Hugh W. Ducklow, Markus B. Karner, Christina M. Preston, and David M. Karl. Abundance and distribution of planktonic *Archaea* and *Bacteria* in the waters west of the Antarctic Peninsula. Limnology and Oceanography, 48(5):1893–1902, September 2003. ISSN 1939-5590. doi: 10.4319/lo.2003.48.5.1893.

R. Eric Collins, Gabrielle Rocap, and Jody W. Deming. Persistence of bacterial and archaeal communities in sea ice through an Arctic winter. Environmental Microbiology, 12(7):1828–1841, July 2010. ISSN 1462-2920. doi: 10.1111/j.1462-2920.2010.02179.x.

G. A. Cutter, K. Casciotti, P. Croot, W. Geibert, L.-E. Heimbürger, M. Lohan, others, and T. van de Flierdt. Sampling and Sample-handling Protocols for GEOTRACES Cruises. Technical report, GEOTRACES, 2017.

James J Davis, Alice R Wattam, Ramy K Aziz, Thomas Brettin, Ralph Butler, Rory M Butler, Philippe Chlenski, Neal Conrad, Allan Dickerman, Emily M Dietrich, Joseph L Gabbard, Svetlana Gerdes, Andrew Guard, Ronald W Kenyon, Dustin Machi, Chunhong Mao, Dan Murphy-Olson, Marcus Nguyen, Eric K Nordberg, Gary J Olsen, Robert D Olson, Jamie C Overbeek, Ross Overbeek, Bruce Parrello, Gordon D Pusch, Maulik Shukla, Chris Thomas, Margo VanOeffelen, Veronika Vonstein, Andrew S Warren, Fangfang Xia, Dawen Xie, Hyunseung Yoo, and Rick Stevens. The PATRIC Bioinformatics Resource Center: expanding data and analysis capabilities. Nucleic Acids Research, October 2019. ISSN 1362-4962. doi: 10.1093/nar/gkz943.

C. Deutsch, N. Gruber, R. M. Key, J. L. Sarmiento, and A. Ganachaud. Denitrification and N_2_ fixation in the Pacific Ocean. Global Biogeochemical Cycles, 15(2):483–506, 2001.

Arwyn Edwards, Karen A. Cameron, Joseph M. Cook, Aliyah R. Debbonaire, Eleanor Furness, Melanie C. Hay, and Sara M.E. Rassner. Microbial genomics amidst the Arctic crisis. Microbial Genomics, 6(5), May 2020. ISSN 2057-5858. doi: 10.1099/mgen.0.000375.

Paul G. Falkowski and Matthew J. Oliver. Mix and match: how climate selects phytoplankton. Nature Reviews Microbiology, 5(10):813–819, October 2007. ISSN 1740-1534. doi: 10.1038/nrmicro1751.

Jesse R. Farmer, Daniel M. Sigman, Julie Granger, Ona M. Underwood, François Fripiat, Thomas M. Cronin, Alfredo Martínez-García, and Gerald H. Haug. Arctic Ocean stratification set by sea level and freshwater inputs since the last ice age. Nature Geoscience, 14(9):684–689, August 2021. ISSN 1752-0908. doi: 10.1038/s41561-021-00789-y.

Clémence Frioux, Dipali Singh, Tamas Korcsmaros, and Falk Hildebrand. From bag-of-genes to bag-of-genomes: metabolic modelling of communities in the era of metagenome-assembled genomes. Computational and Structural Biotechnology Journal, 18:1722–1734, January 2020. ISSN 2001-0370. doi: 10.1016/j.csbj.2020.06.028.

Pierre E. Galand, Carmen Gutiérrez-Provecho, Ramon Massana, Josep M. Gasol, and Emilio O. Casamayor. Inter-annual recurrence of archaeal assemblages in the coastal NW Mediterranean Sea (Blanes Bay Microbial Observatory). Limnology and Oceanography, 55(5):2117–2125, September 2010. ISSN 1939-5590. doi: 10.4319/lo.2010.55.5.2117.

Carlos Garcia-Soto, Lijing Cheng, Levke Caesar, S. Schmidtko, Elizabeth B. Jewett, Alicia Cheripka, Ignatius Rigor, Ainhoa Caballero, Sanae Chiba, Jose Carlos Báez, Tymon Zielinski, and John Patrick Abraham. An Overview of Ocean Climate Change Indicators: Sea Surface Temperature, Ocean Heat Content, Ocean pH, Dissolved Oxygen Concentration, Arctic Sea Ice Extent, Thickness and Volume, Sea Level and Strength of the AMOC (Atlantic Meridional Overturning Circulation). Frontiers in Marine Science, 8, September 2021. ISSN 2296-7745. doi: 10.3389/fmars.2021.642372.

Donglai Gong and Robert S. Pickart. Early summer water mass transformation in the eastern Chukchi Sea. Deep Sea Research Part II: Topical Studies in Oceanography, 130:43–55, August 2016. ISSN 0967-0645. doi: 10.1016/j.dsr2.2016.04.015.

N. Gruber and J. L. Sarmiento. Global patterns of marine nitrogen fixation and denitrification. Global Biogeochemical Cycles, 11(2):235–266, 1997.

Lionel Guy and Thijs J.G. Ettema. The archaeal ‘TACK’ superphylum and the origin of eukaryotes. Trends in Microbiology, 19(12):580–587, December 2011. ISSN 0966-842X. doi: 10.1016/j.tim.2011.09.002.

Jose M Haro-Moreno, Francisco Rodriguez-Valera, Purificación López-García, David Moreira, and Ana-Belen Martin-Cuadrado. New insights into marine group III Euryarchaeota, from dark to light. The ISME Journal, 11(5):1102–1117, January 2017. ISSN 1751-7370. doi: 10.1038/ismej.2016.188.

Lydie Herfort, Stefan Schouten, Ben Abbas, Marcel J. W. Veldhuis, Marco J. L. Coolen, Cornelia Wuchter, Jan P. Boon, Gerhard J. Herndl, and Jaap S. Sinninghe DamstÃ©. Variations in spatial and temporal distribution of Archaea in the North Sea in relation to environmental variables: Marine Archaea in the North Sea. FEMS Microbiology Ecology, 62(3):242–257, December 2007. ISSN 1574-6941. doi: 10.1111/j.1574-6941.2007.00397.x.

Ove Hoegh-Guldberg, Elvira S. Poloczanska, William Skirving, and Sophie Dove. Coral Reef Ecosystems under Climate Change and Ocean Acidification. Frontiers in Marine Science, 4, May 2017. ISSN 2296-7745. doi: 10.3389/fmars.2017.00158.

Doug Hyatt, Gwo-Liang Chen, Philip F LoCascio, Miriam L Land, Frank W Larimer, and Loren J Hauser. Prodigal: prokaryotic gene recognition and translation initiation site identification. BMC Bioinformatics, 11(1), March 2010. ISSN 1471-2105. doi: 10.1186/1471-2105-11-119.

Danny Ionescu, Sigrid Penno, Maya Haimovich, Branko Rihtman, Aram Goodwin, Daniel Schwartz, Lena Hazanov, Mark Chernihovsky, Anton F. Post, and Aharon Oren. Archaea in the Gulf of Aqaba: Archaea in the Gulf of Aqaba. FEMS Microbiology Ecology, 69(3):425–438, September 2009. ISSN 1574-6941. doi: 10.1111/j.1574-6941.2009.00721.x.

L. T. Jensen, N. J. Wyatt, B. S. Twining, S. Rauschenberg, W. M. Landing, R. M. Sherrell, and J. N. Fitzsimmons. Biogeochemical Cycling of Dissolved Zinc in the Western Arctic (Arctic GEOTRACES GN01). Global Biogeochemical Cycles, 33(3):343–369, March 2019. ISSN 1944-9224. doi: 10.1029/2018gb005975.

O.M. Johannessen, L. Bengtsson, M.W. Miles, S. I. Kuzmina, V.A. Semenov, G.V. Alekseev, A.P. Nagurnyi, V.F. Zakharov, L.P. Bobylev, L.H. Pettersson, K. Hasselmann, and H.P. Cattle. Arctic climate change: observed and modelled temperature and sea-ice variability. Tellus A: Dynamic Meteorology and Oceanography, 56(5):559, January 2004. ISSN 1600-0870. doi: 10.3402/tellusa.v56i5.14599.

E. P. Jones and L. G. Anderson. On the origin of the chemical properties of the Arctic Ocean halocline. Journal of Geophysical Research: Oceans, 91(C9):10759–10767, September 1986. ISSN 0148-0227. doi: 10.1029/jc091ic09p10759.

Karen Junge, Christopher Krembs, Jody Deming, Aaron Stierle, and Hajo Eicken. A microscopic approach to investigate bacteria under in situ conditions in sea-ice samples. Annals of Glaciology, 33:304–310, 2001. ISSN 1727-5644. doi: 10.3189/172756401781818275.

D. Kadko and W. Landing. U.S. Arctic GEOTRACES: USCGC Healy (HLY1502). https://www.geotraces.org/gn01/, 2015.

Dongwan D. Kang, Feng Li, Edward Kirton, Ashleigh Thomas, Rob Egan, Hong An, and Zhong Wang. MetaBAT 2: an adaptive binning algorithm for robust and efficient genome reconstruction from metagenome assemblies. PeerJ, 7:e7359, July 2019. ISSN 2167-8359. doi: 10.7717/peerj.7359.

Markus B. Karner, Edward F. DeLong, and David M. Karl. Archaeal dominance in the mesopelagic zone of the Pacific Ocean. Nature, 409(6819):507–510, January 2001. ISSN 1476-4687. doi: 10.1038/35054051.

Lester Kwiatkowski, Olivier Torres, Laurent Bopp, Olivier Aumont, Matthew Chamberlain, James R. Christian, John P. Dunne, Marion Gehlen, Tatiana Ilyina, Jasmin G. John, Andrew Lenton, Hongmei Li, Nicole S. Lovenduski, James C. Orr, Julien Palmieri, Yeray Santana-Falcón, Jörg Schwinger, Roland Séférian, Charles A. Stock, Alessandro Tagliabue, Yohei Takano, Jerry Tjiputra, Katsuya Toyama, Hiroyuki Tsujino, Michio Watanabe, Akitomo Yamamoto, Andrew Yool, and Tilo Ziehn. Twenty-first century ocean warming, acidification, deoxygenation, and upper-ocean nutrient and primary production decline from CMIP6 model projections. Biogeosciences, 17(13): 3439–3470, July 2020. ISSN 1726-4189. doi: 10.5194/bg-17-3439-2020.

Ivica Letunic and Peer Bork. Interactive Tree Of Life (iTOL): an online tool for phylogenetic tree display and annotation. Bioinformatics, 23(1):127–128, October 2006. ISSN 1367-4803. doi: 10.1093/bioinformatics/btl529.

Haodong Liu, Chuanlun L. Zhang, Chunyan Yang, Songze Chen, Zhiwei Cao, Zhiwei Zhang, and Jiwei Tian. Marine Group II Dominates Planktonic Archaea in Water Column of the Northeastern South China Sea. Frontiers in Microbiology, 8, June 2017. ISSN 1664-302X. doi: 10.3389/fmicb.2017.01098.

Qian Liu, Bradley B Tolar, Meredith J Ross, Jelani B Cheek, Corinne M Sweeney, Natalie J Wallsgrove, Brian N Popp, and James T Hollibaugh. Light and temperature control the seasonal distribution of thaumarchaeota in the South Atlantic bight. The ISME Journal, 12(6):1473–1485, February 2018. ISSN 1751-7370. doi: 10.1038/s41396-018-0066-4.

Yong-Xin Liu, Yuan Qin, Tong Chen, Meiping Lu, Xubo Qian, Xiaoxuan Guo, and Yang Bai. A practical guide to amplicon and metagenomic analysis of microbiome data. Protein & Cell, 12(5): 315–330, May 2020. ISSN 1674-8018. doi: 10.1007/s13238-020-00724-8.

Willm Martens-Habbena, Paul M. Berube, Hidetoshi Urakawa, José R. de la Torre, and David A. Stahl. Ammonia oxidation kinetics determine niche separation of nitrifying Archaea and Bacteria. Nature, 461(7266):976–979, September 2009. ISSN 1476-4687. doi: 10.1038/nature08465.

F. A. Matsen, R. B. Kodner, and E. V. Armbrust. pplacer: linear time maximum-likelihood and Bayesian phylogenetic placement of sequences onto a fixed reference tree. BMC Bioinformatics, 11:538, 2010.

Stephanie N. Merbt, David A. Stahl, Emilio O. Casamayor, Eugènia Martí, Graeme W. Nicol, and James I. Prosser. Differential photoinhibition of bacterial and archaeal ammonia oxidation. FEMS Microbiology Letters, 327(1):41–46, December 2011. ISSN 0378-1097. doi: 10.1111/j.1574-6968.2011.02457.x.

Alexandra Meziti, Luis M. Rodriguez-R, Janet K. Hatt, Angela Peña-Gonzalez, Karen Levy, and Konstantinos T. Konstantinidis. The Reliability of Metagenome-Assembled Genomes (MAGs) in Representing Natural Populations: Insights from Comparing MAGs against Isolate Genomes Derived from the Same Fecal Sample. Applied and Environmental Microbiology, 87(6), February 2021. ISSN 1098-5336. doi: 10.1128/aem.02593-20.

Xosé Anxelu G. Morán, Ángel López-Urrutia, Alejandra Calvo-Díaz, and William K. W. LI. Increasing importance of small phytoplankton in a warmer ocean. Global Change Biology, 16(3):1137–1144, January 2010. ISSN 1365-2486. doi: 10.1111/j.1365-2486.2009.01960.x.

Stephen Nayfach, Simon Roux, Rekha Seshadri, Daniel Udwary, Neha Varghese, Frederik Schulz, Dongying Wu, David Paez-Espino, I-Min Chen, Marcel Huntemann, Krishna Palaniappan, Joshua Ladau, Supratim Mukherjee, T. B. K. Reddy, Torben Nielsen, Edward Kirton, José P. Faria, Janaka N. Edirisinghe, Christopher S. Henry, Sean P. Jungbluth, Dylan Chivian, Paramvir Dehal, Elisha M. Wood-Charlson, Adam P. Arkin, Susannah G. Tringe, Axel Visel, Helena Abreu, Silvia G. Acinas, Eric Allen, Michelle A. Allen, Lauren V. Alteio, Gary Andersen, Alexandre M. Anesio, Graeme Attwood, Viridiana Avila-Magaña, Yacine Badis, Jake Bailey, Brett Baker, Petr Baldrian, Hazel A. Barton, David A. C. Beck, Eric D. Becraft, Harry R. Beller, J. Michael Beman, Rizlan Bernier-Latmani, Timothy D. Berry, Anthony Bertagnolli, Stefan Bertilsson, Jennifer M. Bhatnagar, Jordan T. Bird, Jeffrey L. Blanchard, Sara E. Blumer-Schuette, Brendan Bohannan, Mikayla A. Borton, Allyson Brady, Susan H. Brawley, Juliet Brodie, Steven Brown, Jennifer R. Brum, Andreas Brune, Donald A. Bryant, Alison Buchan, Daniel H. Buckley, Joy Buongiorno, Hinsby Cadillo-Quiroz, Sean M. Caffrey, Ashley N. Campbell, Barbara Campbell, Stephanie Carr, JoLynn Carroll, S. Craig Cary, Anna M. Cates, Rose Ann Cattolico, Ricardo Cavicchioli, Ludmila Chistoserdova, Maureen L. Coleman, Philippe Constant, Jonathan M. Conway, Walter P. Mac Cormack, Sean Crowe, Byron Crump, Cameron Currie, Rebecca Daly, Kristen M. DeAngelis, Vincent Denef, Stuart E. Denman, Adey Desta, Hebe Dionisi, Jeremy Dodsworth, Nina Dombrowski, Timothy Donohue, Mark Dopson, Timothy Driscoll, Peter Dunfield, Christopher L. Dupont, Katherine A. Dynarski, Virginia Edgcomb, Elizabeth A. Edwards, Mostafa S. Elshahed, Israel Figueroa, Beverly Flood, Nathaniel Fortney, Caroline S. Fortunato, Christopher Francis, Claire M. M. Gachon, Sarahi L. Garcia, Maria C. Gazitua, Terry Gentry, Lena Gerwick, Javad Gharechahi, Peter Girguis, John Gladden, Mary Gradoville, Stephen E. Grasby, Kelly Gravuer, Christen L. Grettenberger, Robert J. Gruninger, Jiarong Guo, Mussie Y. Habteselassie, Steven J. Hallam, Roland Hatzenpichler, Bela Hausmann, Terry C. Hazen, Brian Hedlund, Cynthia Henny, Lydie Herfort, Maria Hernandez, Olivia S. Hershey, Matthias Hess, Emily B. Hollister, Laura A. Hug, Dana Hunt, Janet Jansson, Jessica Jarett, Vitaly V. Kadnikov, Charlene Kelly, Robert Kelly, William Kelly, Cheryl A. Kerfeld, Jeff Kimbrel, Jonathan L. Klassen, Konstantinos T. Konstantinidis, Laura L. Lee, Wen-Jun Li, Andrew J. Loder, Alexander Loy, Mariana Lozada, Barbara MacGregor, Cara Magnabosco, Aline Maria da Silva, R. Michael McKay, Katherine McMahon, Chris S. McSweeney, Mónica Medina, Laura Meredith, Jessica Mizzi, Thomas Mock, Lily Momper, Mary Ann Moran, Connor Morgan-Lang, Duane Moser, Gerard Muyzer, David Myrold, Maisie Nash, Camilla L. Nesbø, Anthony P. Neumann, Rebecca B. Neumann, Daniel Noguera, Trent Northen, Jeanette Norton, Brent Nowinski, Klaus Nüsslein, Michelle A. O’Malley, Rafael S. Oliveira, Valeria Maia de Oliveira, Tullis Onstott, Jay Osvatic, Yang Ouyang, Maria Pachiadaki, Jacob Parnell, Laila P. Partida-Martinez, Kabir G. Peay, Dale Pelletier, Xuefeng Peng, Michael Pester, Jennifer Pett-Ridge, Sari Peura, Petra Pjevac, Alvaro M. Plominsky, Anja Poehlein, Phillip B. Pope, Nikolai Ravin, Molly C. Redmond, Rebecca Reiss, Virginia Rich, Christian Rinke, Jorge L. Mazza Rodrigues, William Rodriguez-Reillo, Karen Rossmassler, Joshua Sackett, Ghasem Hosseini Salekdeh, Scott Saleska, Matthew Scarborough, Daniel Schachtman, Christopher W. Schadt, Matthew Schrenk, Alexander Sczyrba, Aditi Sengupta, Joao C. Setubal, Ashley Shade, Christine Sharp, David H. Sherman, Olga V. Shubenkova, Isabel Natalia Sierra-Garcia, Rachel Simister, Holly Simon, Sara Sjöling, Joan Slonczewski, Rafael Soares Correa de Souza, John R. Spear, James C. Stegen, Ramunas Stepanauskas, Frank Stewart, Garret Suen, Matthew Sullivan, Dawn Sumner, Brandon K. Swan, Wesley Swingley, Jonathan Tarn, Gordon T. Taylor, Hanno Teeling, Memory Tekere, Andreas Teske, Torsten Thomas, Cameron Thrash, James Tiedje, Claire S. Ting, Benjamin Tully, Gene Tyson, Osvlado Ulloa, David L. Valentine, Marc W. Van Goethem, Jean VanderGheynst, Tobin J. Verbeke, John Vollmers, Aurèle Vuillemin, Nicholas B. Waldo, David A. Walsh, Bart C. Weimer, Thea Whitman, Paul van der Wielen, Michael Wilkins, Timothy J. Williams, Ben Woodcroft, Jamie Woolet, Kelly Wrighton, Jun Ye, Erica B. Young, Noha H. Youssef, Feiqiao Brian Yu, Tamara I. Zemskaya, Ryan Ziels, Tanja Woyke, Nigel J. Mouncey, Natalia N. Ivanova, Nikos C. Kyrpides, and Emiley A. Eloe-Fadrosh. A genomic catalog of Earth’s microbiomes. Nature Biotechnology, 39(4):499–509, November 2020. ISSN 1546-1696. doi: 10.1038/s41587-020-0718-6.

Sergey Nurk, Dmitry Meleshko, Anton Korobeynikov, and Pavel A. Pevzner. metaSPAdes: a new versatile metagenomic assembler. Genome Research, 27(5):824–834, March 2017. ISSN 1549-5469. doi: 10.1101/gr.213959.116.

Pierre Offre, Anja Spang, and Christa Schleper. Archaea in Biogeochemical Cycles. Annual Review of Microbiology, 67(1):437–457, September 2013. ISSN 1545-3251. doi: 10.1146/annurev-micro-092412-155614.

J. Oksanen, G. L. Simpson, F. G. Blanchet, R. Kindt, P. Legendre, P. R. Minchin, R. B. O’Hara, P. Solymos, M. H. H. Stevens, E. Szoecs, H. Wagner, et al. vegan: Community Ecology Package, 2022. R package version 2.6-2.

L. Oziel, V. Schourup-Kristensen, C. Wekerle, and J. Hauck. The Pan-Arctic Continental Slope as an Intensifying Conveyer Belt for Nutrients in the Central Arctic Ocean (1985–2015). Global Biogeochemical Cycles, 36(6), June 2022. ISSN 1944-9224. doi: 10.1029/2021gb007268.

Mary I. O’Connor, Michael F. Piehler, Dina M. Leech, Andrea Anton, and John F. Bruno. Warming and Resource Availability Shift Food Web Structure and Metabolism. PLoS Biology, 7(8):e1000178, August 2009. ISSN 1545-7885. doi: 10.1371/journal.pbio.1000178.

Donovan H. Parks, Michael Imelfort, Connor T. Skennerton, Philip Hugenholtz, and Gene W. Tyson. CheckM: assessing the quality of microbial genomes recovered from isolates, single cells, and metagenomes. Genome Research, 25(7):1043–1055, May 2015. ISSN 1549-5469. doi: 10.1101/gr.186072.114.

Olivier Pereira, Corentin Hochart, Jean Christophe Auguet, Didier Debroas, and Pierre E. Galand. Genomic ecology of Marine Group II, the most common marine planktonic Archaea across the surface ocean. MicrobiologyOpen, 8(9), July 2019. ISSN 2045-8827. doi: 10.1002/mbo3.852.

Bruce J. Peterson, James McClelland, Ruth Curry, Robert M. Holmes, John E. Walsh, and Knut Aagaard. Trajectory Shifts in the Arctic and Subarctic Freshwater Cycle. Science, 313(5790): 1061–1066, August 2006. ISSN 1095-9203. doi: 10.1126/science.1122593.

Maria N. Pisareva, Robert S. Pickart, M.A. Spall, C. Nobre, D.J. Torres, G.W.K. Moore, and Terry E. Whitledge. Flow of pacific water in the western Chukchi Sea: Results from the 2009 RUSALCA expedition. Deep Sea Research Part I: Oceanographic Research Papers, 105:53–73, November 2015. ISSN 0967-0637. doi: 10.1016/j.dsr.2015.08.011.

Angela Pitcher, Cornelia Wuchter, Kathi Siedenberg, Stefan Schouten, and Jaap S. Sinninghe Damsté. Crenarchaeol tracks winter blooms of ammonia-oxidizing Thaumarchaeota in the coastal North Sea. Limnology and Oceanography, 56(6):2308–2318, November 2011. ISSN 1939-5590. doi: 10.4319/lo.2011.56.6.2308.

Wei Qin, Shady A. Amin, Willm Martens-Habbena, Christopher B. Walker, Hidetoshi Urakawa, Allan H. Devol, Anitra E. Ingalls, James W. Moffett, E. Virginia Armbrust, and David A. Stahl. Marine ammonia-oxidizing archaeal isolates display obligate mixotrophy and wide ecotypic variation. Proceedings of the National Academy of Sciences, 111(34):12504–12509, August 2014. ISSN 1091-6490. doi: 10.1073/pnas.1324115111.

R Core Team. R: A Language and Environment for Statistical Computing. R Foundation for Statistical Computing, Vienna, Austria, 2021.

Mika Rantanen, Alexey Yu. Karpechko, Antti Lipponen, Kalle Nordling, Otto Hyvärinen, Kimmo Ruosteenoja, Timo Vihma, and Ari Laaksonen. The Arctic has warmed nearly four times faster than the globe since 1979. Communications Earth & Environment, 3(1), August 2022. ISSN 2662-4435. doi: 10.1038/s43247-022-00498-3.

A. C. Redfield, B. H. Ketchum, and F. A. Richards. The influence of organisms on the composition of sea-water. In M. N. Hill, editor, The Sea: Ideas and Observations on Progress in the Study of the Seas, Vol. 2: The Composition of Sea-Water, Comparative and Descriptive Oceanography, pages 26–77. Interscience, New York, 1963.

Linta Reji and Christopher A Francis. Metagenome-assembled genomes reveal unique metabolic adaptations of a basal marine Thaumarchaeota lineage. The ISME Journal, 14(8):2105–2115, May 2020. ISSN 1751-7370. doi: 10.1038/s41396-020-0675-6.

Linta Reji, Bradley B Tolar, Jason M Smith, Francisco P Chavez, and Christopher A Francis. Differential co-occurrence relationships shaping ecotype diversification within *Thaumarchaeota* populations in the coastal ocean water column. The ISME Journal, 13(5):1144–1158, January 2019. ISSN 1751-7370. doi: 10.1038/s41396-018-0311-x.

Christian Rinke, Patrick Schwientek, Alexander Sczyrba, Natalia N. Ivanova, Iain J. Anderson, Jan-Fang Cheng, Aaron Darling, Stephanie Malfatti, Brandon K. Swan, Esther A. Gies, Jeremy A. Dodsworth, Brian P. Hedlund, George Tsiamis, Stefan M. Sievert, Wen-Tso Liu, Jonathan A. Eisen, Steven J. Hallam, Nikos C. Kyrpides, Ramunas Stepanauskas, Edward M. Rubin, Philip Hugenholtz, and Tanja Woyke. Insights into the phylogeny and coding potential of microbial dark matter. Nature, 499(7459):431–437, July 2013. ISSN 1476-4687. doi: 10.1038/nature12352.

Christian Rinke, Francesco Rubino, Lauren F Messer, Noha Youssef, Donovan H Parks, Maria Chuvochina, Mark Brown, Thomas Jeffries, Gene W Tyson, Justin R Seymour, and Philip Hugenholtz. A phylogenomic and ecological analysis of the globally abundant Marine Group II archaea (*Ca*. Poseidoniales ord. nov.). The ISME Journal, 13(3):663–675, October 2019. ISSN 1751-7370. doi: 10.1038/s41396-018-0282-y.

Christian Rinke, Maria Chuvochina, Aaron J. Mussig, Pierre-Alain Chaumeil, Adrian A. Davin, David W. Waite, William B Whitman, Donovan H. Parks, and Philip Hugenholtz. Resolving widespread incomplete and uneven archaeal classifications based on a rank-normalized genome-based taxonomy. bioRxiv, March 2020. doi: 10.1101/2020.03.01.972265.

Anindya Roy Chowdhury and Chitra Dutta. A pursuit of lineage-specific and niche-specific proteome features in the world of archaea. BMC Genomics, 13(1):236, 2012. ISSN 1471-2164. doi: 10.1186/1471-2164-13-236.

Bert Rudels. Arctic Ocean circulation, processes and water masses: A description of observations and ideas with focus on the period prior to the International Polar Year 2007–2009. Progress in Oceanography, 132:22–67, March 2015. ISSN 0079-6611. doi: 10.1016/j.pocean.2013.11.006.

Alyson E. Santoro, Carolyn Buchwald, Matthew R. McIlvin, and Karen L. Casciotti. Isotopic Signature of N _2_ O Produced by Marine Ammonia-Oxidizing Archaea. Science, 333(6047): 1282–1285, September 2011. ISSN 1095-9203. doi: 10.1126/science.1208239.

Reiner Schlitzer. Ocean Data View. https://odv.awi.de, 2023.

Sunke Schmidtko, Lothar Stramma, and Martin Visbeck. Decline in global oceanic oxygen content during the past five decades. Nature, 542(7641):335–339, February 2017. ISSN 1476-4687. doi: 10.1038/nature21399.

Christian M. K. Sieber, Alexander J. Probst, Allison Sharrar, Brian C. Thomas, Matthias Hess, Susannah G. Tringe, and Jillian F. Banfield. Recovery of genomes from metagenomes via a dereplication, aggregation and scoring strategy. Nature Microbiology, 3(7):836–843, May 2018. ISSN 2058-5276. doi: 10.1038/s41564-018-0171-1.

Anja Spang, Jimmy H. Saw, Steffen L. Jørgensen, Katarzyna Zaremba-Niedzwiedzka, Joran Martijn, Anders E. Lind, Roel van Eijk, Christa Schleper, Lionel Guy, and Thijs J. G. Ettema. Complex archaea that bridge the gap between prokaryotes and eukaryotes. Nature, 521(7551):173–179, May 2015. ISSN 1476-4687. doi: 10.1038/nature14447.

Alexandros Stamatakis. RAxML-VI-HPC: maximum likelihood-based phylogenetic analyses with thousands of taxa and mixed models. Bioinformatics, 22(21):2688–2690, August 2006. ISSN 1367-4803. doi: 10.1093/bioinformatics/btl446.

H. Teeling and F. O. Glockner. Current opportunities and challenges in microbial metagenome analysis–a bioinformatic perspective. Briefings in Bioinformatics, 13(6):728–742, September 2012. ISSN 1477-4054. doi: 10.1093/bib/bbs039.

Ziye Wang, Pingqin Huang, Ronghui You, Fengzhu Sun, and Shanfeng Zhu. MetaBinner: a high-performance and stand-alone ensemble binning method to recover individual genomes from complex microbial communities. Genome Biology, 24(1), January 2023. ISSN 1474-760X. doi: 10.1186/s13059-022-02832-6.

Laura M. Whitmore, Peter L. Morton, Benjamin S. Twining, and Alan M. Shiller. Vanadium cycling in the Western Arctic Ocean is influenced by shelf-basin connectivity. Marine Chemistry, 216: 103701, October 2019. ISSN 0304-4203. doi: 10.1016/j.marchem.2019.103701.

Laura M. Whitmore, Alan M. Shiller, Tristan J. Horner, Yang Xiang, Maureen E. Auro, Dorothea Bauch, Frank Dehairs, Phoebe J. Lam, Jingxuan Li, Maria T. Maldonado, Chantal Mears, Robert Newton, Angelica Pasqualini, Hélène Planquette, Robert Rember, and Helmuth Thomas. Strong Margin Influence on the Arctic Ocean Barium Cycle Revealed by Pan-Arctic Synthesis. Journal of Geophysical Research: Oceans, 127(4), March 2022. ISSN 2169-9291. doi: 10.1029/2021jc017417.

Carl R. Woese and George E. Fox. Phylogenetic structure of the prokaryotic domain: The primary kingdoms. Proceedings of the National Academy of Sciences, 74(11):5088–5090, November 1977. ISSN 1091-6490. doi: 10.1073/pnas.74.11.5088.

R. A. Woodgate, K. Aagaard, R. D. Muench, J. Gunn, G. Björk, B. Rudels, others, and U. Schauer. The Arctic Ocean Boundary Current along the Eurasian slope and the adjacent Lomonosov Ridge: Water mass properties, transports and transformations from moored instruments. Deep-Sea Research Part I: Oceanographic Research Papers, 48:1757–1792, 2001.

Yu-Wei Wu, Blake A. Simmons, and Steven W. Singer. MaxBin 2.0: an automated binning algorithm to recover genomes from multiple metagenomic datasets. Bioinformatics, 32(4):605–607, October 2015. ISSN 1367-4803. doi: 10.1093/bioinformatics/btv638.

Cornelia Wuchter, Ben Abbas, Marco J. L. Coolen, Lydie Herfort, Judith van Bleijswijk, Peer Timmers, Marc Strous, Eva Teira, Gerhard J. Herndl, Jack J. Middelburg, Stefan Schouten, and Jaap S. Sinninghe Damsté. Archaeal nitrification in the ocean. Proceedings of the National Academy of Sciences, 103(33):12317–12322, August 2006. ISSN 1091-6490. doi: 10.1073/pnas.0600756103.

Pelin Yilmaz, Pablo Yarza, Josephine Z. Rapp, and Frank O. Glöckner. Expanding the World of Marine Bacterial and Archaeal Clades. Frontiers in Microbiology, 6, January 2016. ISSN 1664-302X. doi: 10.3389/fmicb.2015.01524.

